# Chemotherapeutic regulation of the ROS/MondoA-dependent TXNIP/GDF15 axis; and derivation of a new organoid metric as a predictive biomarker

**DOI:** 10.1101/2023.08.10.552749

**Authors:** Jinhai Deng, Teng Pan, Yourae Hong, Zaoqu Liu, Xingang Zhou, Zhengwen An, Lifeng Li, Giovanna Alfano, Gang Li, Luigi Dolcetti, Rachel Evans, Jose M Vicencio, Petra Vlckova, Yue Chen, James Monypenny, Camila Araujo De Carvalho Gomes, Kenrick Ng, Caitlin McCarthy, Xiaoping Yang, Zedong Hu, Joanna C. Porter, Christopher J Tape, Mingzhu Yin, Manuel Rodriguez-Justo, Sabine Tejpar, Richard Beatson, Tony Ng

## Abstract

Chemotherapy, the standard of care treatment for cancer patients with advanced disease, has been increasingly recognised to activate host immune responses to produce durable outcomes. Here, in colorectal adenocarcinoma (CRC) we identify chemotherapy-induced Thioredoxin Interacting Protein (*TXNIP),* a MondoA-dependent tumor suppressor gene, as a negative regulator of Growth/Differentiation Factor 15 (GDF15). GDF15 is a negative prognostic factor in CRC and promotes the differentiation of regulatory T cells (Tregs), through CD48 ligation. Intriguingly, multiple models including patient-derived tumor organoids demonstrate that loss of TXNIP/GDF15 axis functionality is associated with advanced disease or chemotherapeutic resistance, with transcriptomic or proteomic GDF15/TXNIP ratios showing potential as a prognostic biomarker. These findings illustrate a potentially common pathway where chemotherapy-induced epithelial stress drives local immune remodelling for patient benefit, with disruption of this pathway seen in refractory or advanced cases.

## Introduction

Colorectal adenocarcinoma (CRC) has the fourth highest mortality amongst cancers, and is characterized by its aggression and heterogeneity^1,2^. Randomized controlled clinical trials have established that chemotherapy results in improved clinical outcomes^3^. 5-FU (fluorouracil), oxaliplatin and irinotecan are the foundation of first-line (FOLFOX) and second-line (FOLFIRI)^4^ treatment respectively. Despite mechanistic differences, all chemotherapy regimens induce apoptosis of replicating cells, leading to a reduction in tumor volume. Chemotherapeutic regimens have historically been regarded as immunologically silent or toxic, however, this view is being increasingly challenged with reports showing that these treatments can modulate immune cells within the tumor microenvironment (TME)^5,6^.

Harnessing the immune system is crucial in achieving long-term durability of response^7^, and chemotherapy reportedly activates anti-tumor immune responses through several mechanisms^8–12^. For example, chemotherapy-induced immunogenic cell death (ICD) leads to cells exposing or releasing damage-associated molecular patterns (DAMPs), such as HSP70, calreticulin, ATP, high-mobility group box 1 (HMGB1), type I IFN, cancer cell-derived nucleic acids and annexin A1^9,10^. These mediators drive anti-tumor immune responses via innate immune cells (dendritic cells [DC], macrophages, NK cells, γδT cells) and adaptive immune cells (T and B cells). Additionally, chemotherapy has been shown to upregulate HLA expression and alter the peptides presented on MHC class I molecules, enabling an antitumor T cell response through the expression of, and reaction to, neo-antigens^8^. Other chemotherapy-induced anti-tumor immunological mechanisms include the down-regulation of immune checkpoint molecules (e.g. PD-L1)^11,12^, however, knowledge of these mechanisms has not yet been translated into a targeted chemo-immunotherapeutic treatment regimes. These anti-tumor immunological benefits of chemotherapy are, of course, balanced by pro-tumor impacts; chemotherapy-induced apoptosis itself, whether epithelial or immune, has been shown to be associated with immunosuppression in multiple cancers^13,14^.

Thioredoxin-interacting protein (TXNIP), an alpha-arrestin protein, is commonly considered a master regulator of cellular oxidation, regulating the expression of Thioredoxin (Trx) via direct binding^15,16^. It has been seen to be silenced by genetic or epigenetic events in a wide range of human tumors, whilst TXNIP-deficient mice have a higher incidence of spontaneous hepatocellular carcinoma^17–20^. Consequently, TXNIP is considered a tumor suppressor gene (TSG). In cell biology, TXNIP has been reported to regulate the cell cycle, oxidative stress responses, angiogenesis, glycolysis and the NLRP3 inflammasome^21–29^. Previous studies have shown chemotherapy drives an increase of TXNIP expression leading to cell cycle arrest and death in epithelial cells^30,31^, however, there are currently no studies that assess the effect of chemotherapy-induced TXNIP expression on the cells that survive chemotherapy, and an understanding of their impact on the TME.

Growth/Differentiation Factor 15 (GDF15), is a distant member of the TGF-β superfamily^32^. At rest, GDF15 is produced at low levels by most epithelial tissues, however in cancers it is frequently over-expressed, particularly in hepatocellular carcinoma, prostate cancer and colorectal cancer^33,34^. Initially, GDF15 was identified as anti-tumorigenic protein with pro-apoptotic capability^35^. However, its tumor-promoting effects are now well-documented to the extent that it is being promoted as a serological biomarker, with increased concentrations being associated with progression, recurrence and death^36,37^, whilst over-expressing or knock-out murine models have demonstrated its promotional role in tumorigenesis^38^. Immunologically, GDF15 is considered an anti-inflammatory factor, supported by the evidence that ubiquitous overexpression decreased systemic inflammatory responses^39^ alongside its negative functional impact on macrophages, dendritic cells and NK cells, coupled with its ability to induce Tregs^40–42^. As a soluble protein, GDF15 exerts its effects by binding to its cognate receptors. To date, there are three receptors reported: Transforming Growth Factor-beta receptor II (TGF-βRII), GDNF-family receptor a-like (GFRAL) and CD48 receptor (SLAMF2).

In this work, using a variety of *in vitro* models, including patient-derived tumor organoids (PDTOs) we demonstrate that oxaliplatin-based chemotherapy reshapes the TME via an increase in ROS-mediated MondoA-dependent TXNIP expression, resulting in decreased expression and secretion of GDF15, leading to a decrease in regulatory T cell (Treg) differentiation. To support the concept of a TXNIP/GDF15/Treg regulatory axis *in situ,* an anti-correlation of TXNIP and GDF15 was observed in matched fresh patient tissue (pre and post chemotherapy), fixed tissue, whole tumor transcripts, and single cell seq data, whilst GDF15 was further seen to correlate with Foxp3 in fixed samples and a transcriptomic dataset. With regards clinical impact, both low TXNIP and high GDF15 were shown to be poor prognostic indicators when assessing protein or transcript expression, allowing us to postulate that the inversion of this phenotype through chemotherapeutic treatment may be associated with positive outcome. Further to this, the axis was seen to be unresponsive in CRC cell lines derived from secondary sites, in a similar manner to chemotherapy-resistant CRC cell lines, with aggressive primary tumours also showing a similar trend. These data suggest that the loss of a responsive axis allows for tumor survival, with this concept being supported by transcriptomic analysis of primary and metastatic disease and responsive and non-responsive cases. Beyond the biology, this study illustrates the potential of the pre-treatment GDF15/TXNIP ratio as a tool to predict chemotherapeutic response in patients allowing for appropriate immunotherapy (GDF15 antagonists in this case) to be administered to non-responders at an early timepoint in a precise and informed manner.

## Results

### TXNIP is upregulated after chemotherapy and associated with favourable prognosis

TXNIP is relatively well-studied in cancer and has been reported to have tumor-suppressive effects as discussed^43^. In CRC, TXNIP expression has been observed to be decreased in tumor cases compared to normal tissues^44^. In support of this, analysis of the TCGA COAD (CRC) database showed decreased *TXNIP* mRNA in tumor samples compared to normal controls (Figure S1A). To validate this, we collected 42 CRC patient samples and observed that tumors presented lower expression of TXNIP as compared to adjacent normal tissue (ANT) (Figure 1A, B). We then used single cell transcriptomics to confirm the same observation in epithelial cells in CRC (Figure 1C).

**Figure 1.**
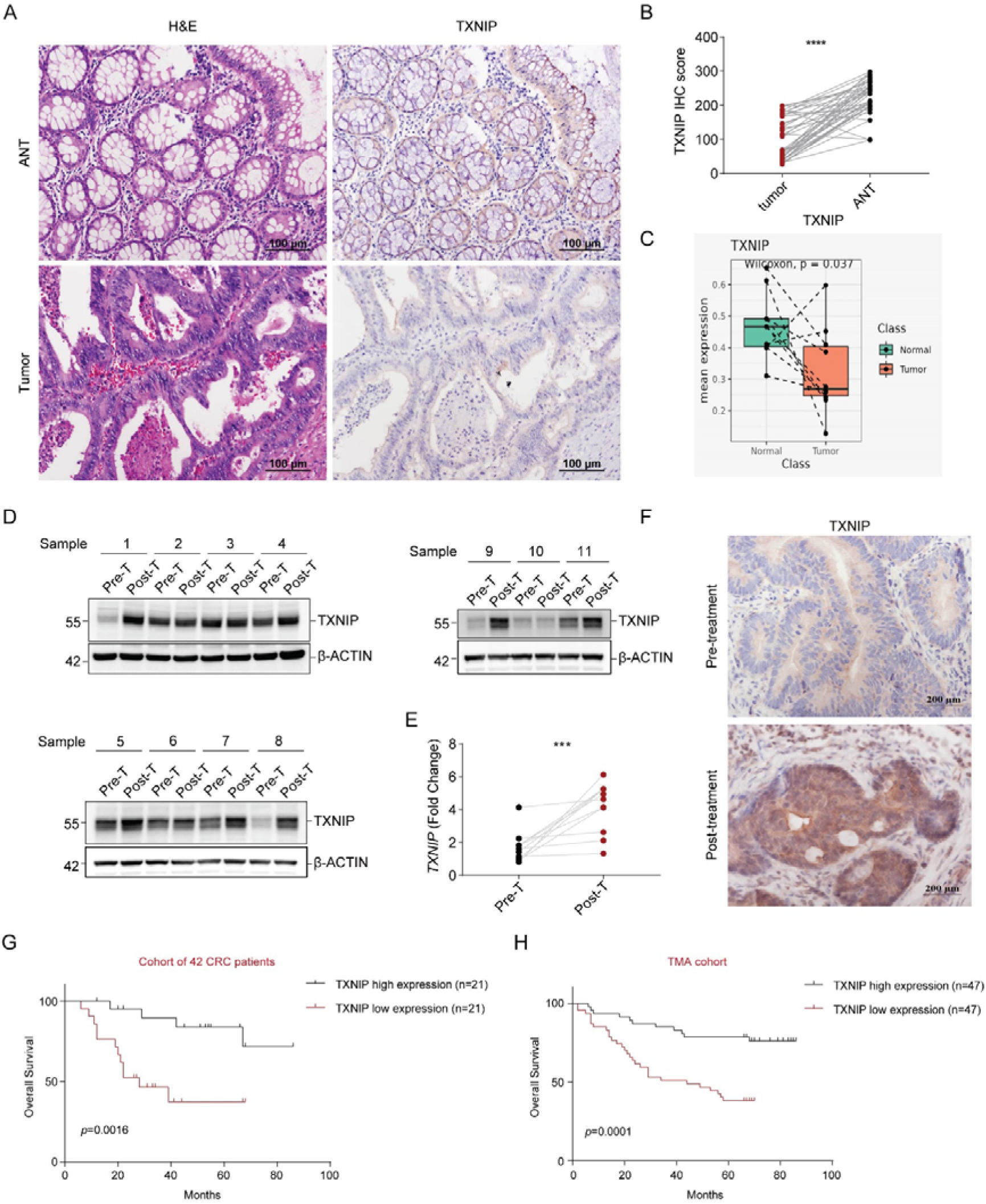
Lower TXNIP expression is observed in CRC tumor samples, however it is increased post-chemotherapeutic treatment. Low levels of TXNIP are associated with poor prognosis. (A) Example of H&E and TXNIP staining in primary CRC tumors and matched adjacent normal tissue (ANT) samples. Magnification ×200. (B) Pooled TXNIP scoring from primary CRC tumors and matched ANT samples (n=42). (C) TXNIP transcript expression in single epithelial cells derived from matched primary CRC tumors and adjacent normal colon (n=10 pairs). (D) TXNIP expression in 11 paired treatment-naïve (Pre-T) tumor samples and oxaliplatin-based neo-adjuvant chemotherapy treated tumor samples (Post-T). (E) TXNIP mRNA levels in samples from D. (F) Example of TXNIP staining in matched Pre-T and Post-T samples. Magnification ×400. (G-H) Kaplan–Meier analysis of overall survival in CRC patients with different TXNIP staining scores from a cohort of 42 CRC patients (G) and CRC tumor tissue microarray (n=94) (H). Data in (G) and (H) were analyzed using two-tailed log-rank test; data in (B) and (E) were analyzed using two-tailed, two-sample unpaired Student’s t test. Data in (C) were analysed using Wilcoxon paired test. Values were expressed as mean ± SEM. ***p < 0.001, ****p < 0.0001, vs. Control

TXNIP has previously been shown to be increased during chemotherapy-induced cell death^30,31^. As TXNIP is considered vital in the regulation of intracellular reactive oxygen species (ROS), which are generated by chemotherapeutic treatment, we questioned whether TXNIP could additionally act as a survival factor. To test this, we took biopsies from CRC patients before and after oxaliplatin-based chemotherapy and measured TXNIP, finding an increase in expression after chemotherapy in 8/11 patients (Figure 1D-F). Somewhat presciently, in light of subsequent findings, 3/11 patients (patients 2, 3 and 10), with advanced disease, showed no increase after treatment (Figure 1D-F). To assess for any association between TXNIP expression and disease progression, and to test whether the chemotherapy inspired increase we had observed would be of benefit to the patient, we used two historic tissue cohorts and two publicly available transcriptomic datasets. High levels of both the protein and the transcript were seen to be associated with favourable prognosis (Figure 1G,H and S1B,C). Moreover, in historic patient cohorts, TXNIP expression was observed to be significantly negatively correlated with clinical stage and lymph node metastasis, with no correlation with respect to age, sex, or tumor size (Table 1 and Table 2)

**Table 1:**
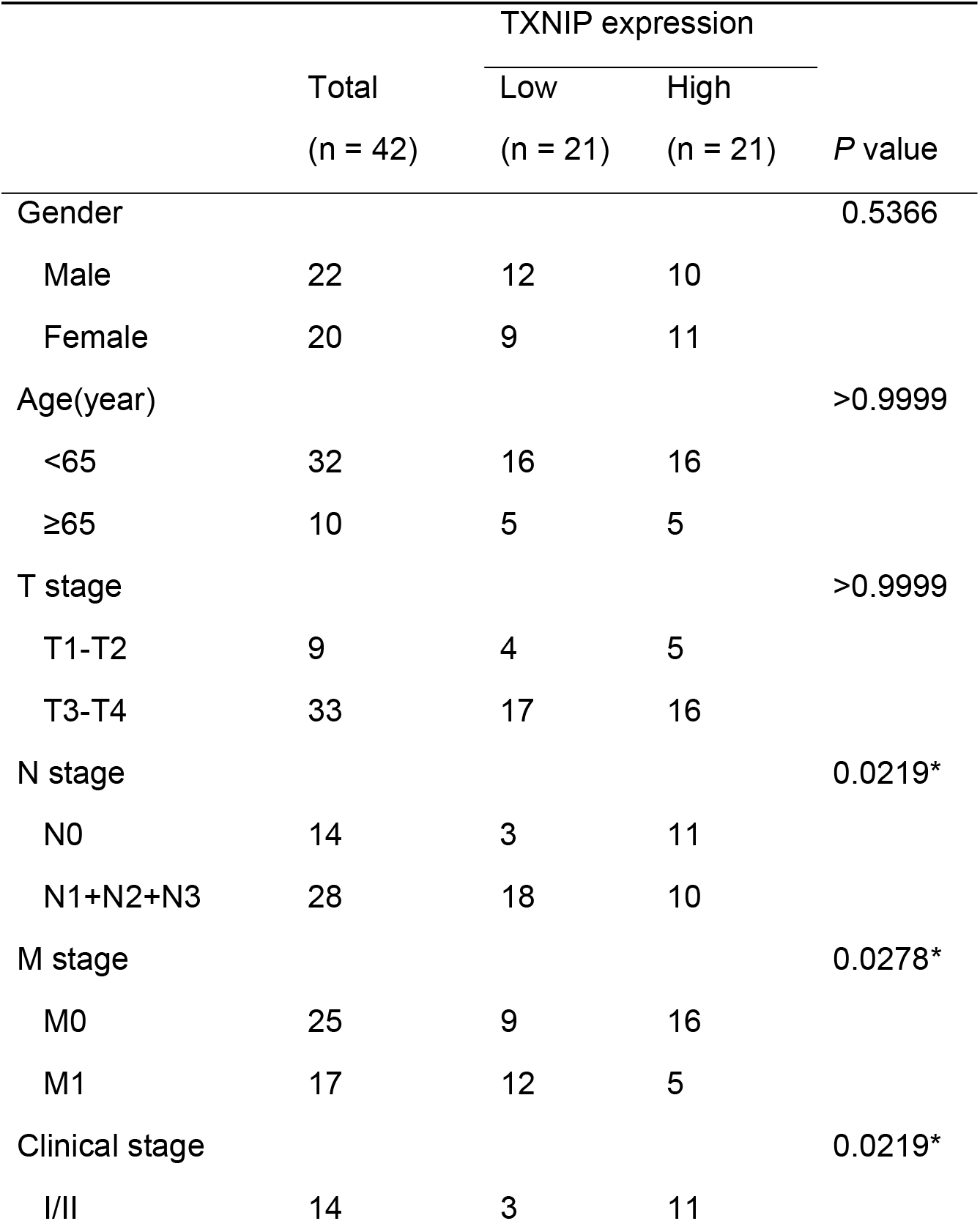
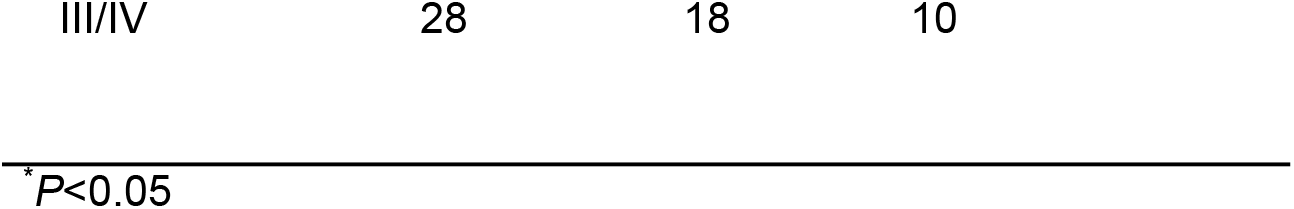
Association between TXNIP expression and clinicopathological features of patients with colorectal cancer in the cohort of 42 CRC patients.

|  | Total<br>(n = 42) | TXNIP expression |  | P value |
| --- | --- | --- | --- | --- |
|  |  | Low<br>(n = 21) | High<br>(n = 21) |  |
| Gender |  |  |  | 0.5366 |
| Male | 22 | 12 | 10 |  |
| Female | 20 | 9 | 11 |  |
| Age(year) |  |  |  | >0.9999 |
| <65 | 32 | 16 | 16 |  |
| ≥65 | 10 | 5 | 5 |  |
| T stage |  |  |  | >0.9999 |
| T1-T2 | 9 | 4 | 5 |  |
| T3-T4 | 33 | 17 | 16 |  |
| N stage |  |  |  | 0.0219* |
| N0 | 14 | 3 | 11 |  |
| N1+N2+N3 | 28 | 18 | 10 |  |
| M stage |  |  |  | 0.0278* |
| M0 | 25 | 9 | 16 |  |
| M1 | 17 | 12 | 5 |  |
| Clinical stage |  |  |  | 0.0219* |
| I/II | 14 | 3 | 11 |  |

|  |  |  |  |
| --- | --- | --- | --- |
| III/IV | 28 | 18 | 10 |

\**P*<0.05

**Table 2:**
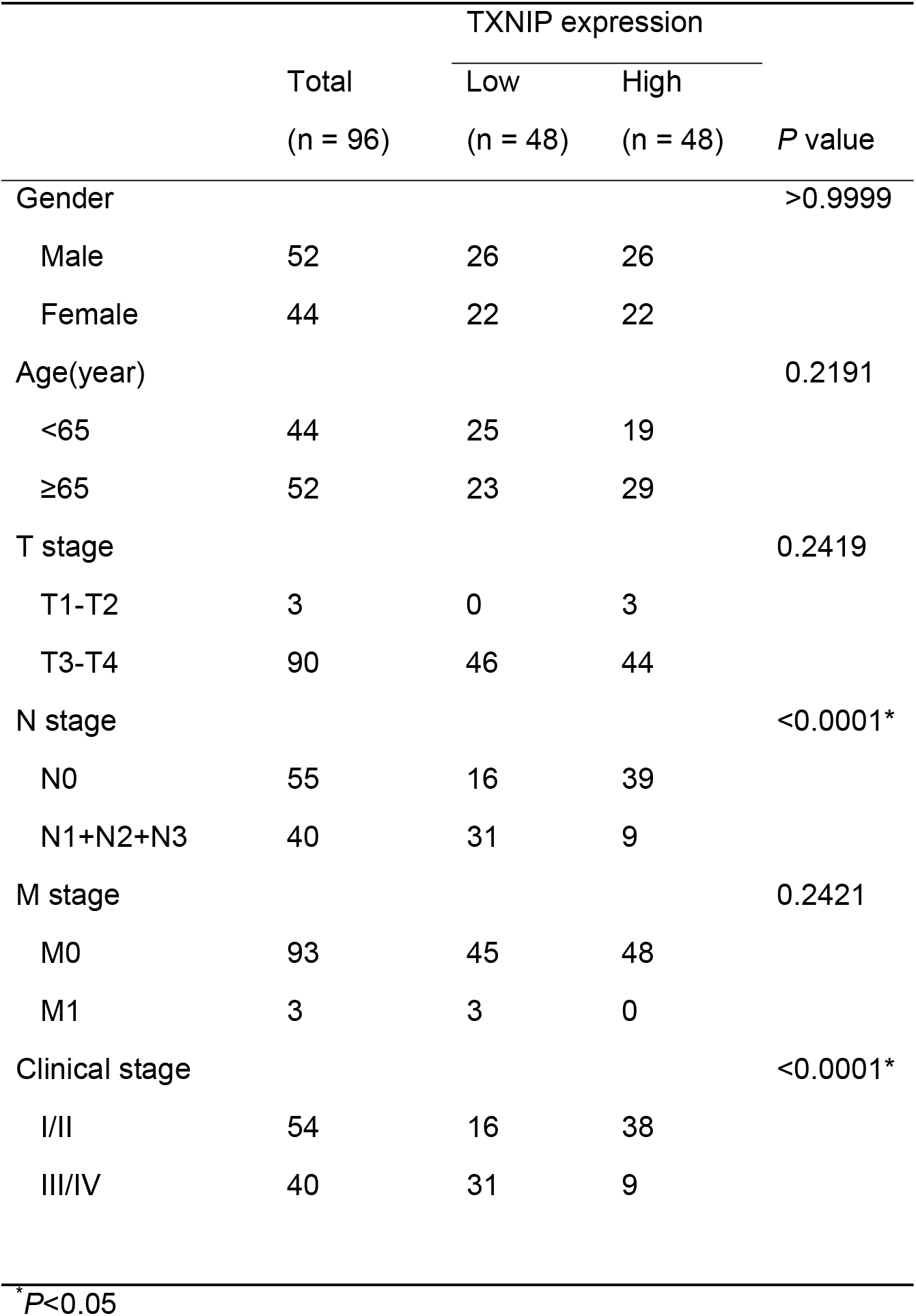
Association between TXNIP expression and clinicopathological features of patients with colorectal cancer in the TMA cohort.

### MondoA regulates chemotherapy-induced TXNIP expression

To assess for the relative expression change of TXNIP after chemotherapy, compared to other transcripts, we used primary colorectal cancer cell lines (DLD1 and HCT15) and treated them with a clinically relevant concentration (10µM)^45^ of oxaliplatin or vehicle. The dead cells were discarded and the live cells were sent for RNA sequencing analysis. The results showed that TXNIP was upregulated as one of the top differentials in both cell lines (Figure 2A, B, Suppl Table 1); validated by RT-PCR and Western blot (Figure 2C-F, S2A-D). Further to this, oxaliplatin upregulated *TXNIP* in a time-dependent (Figure 2C-D, S2A-B) and dose-dependent fashion (Figure 2E-F, S2C-D). 3D (three-dimensional) cell models are reported to be more accurate in mimicking *in vivo* features such as drug responses^46^, therefore we assessed whether this response was observed in cell line-derived spheroids and two patient-derived organoids. In both models we observed the upregulation of TXNIP mRNA (Suppl Fig 2E-H) and protein (Suppl Fig 2I-L) after oxaliplatin treatment.

**Figure 2.**
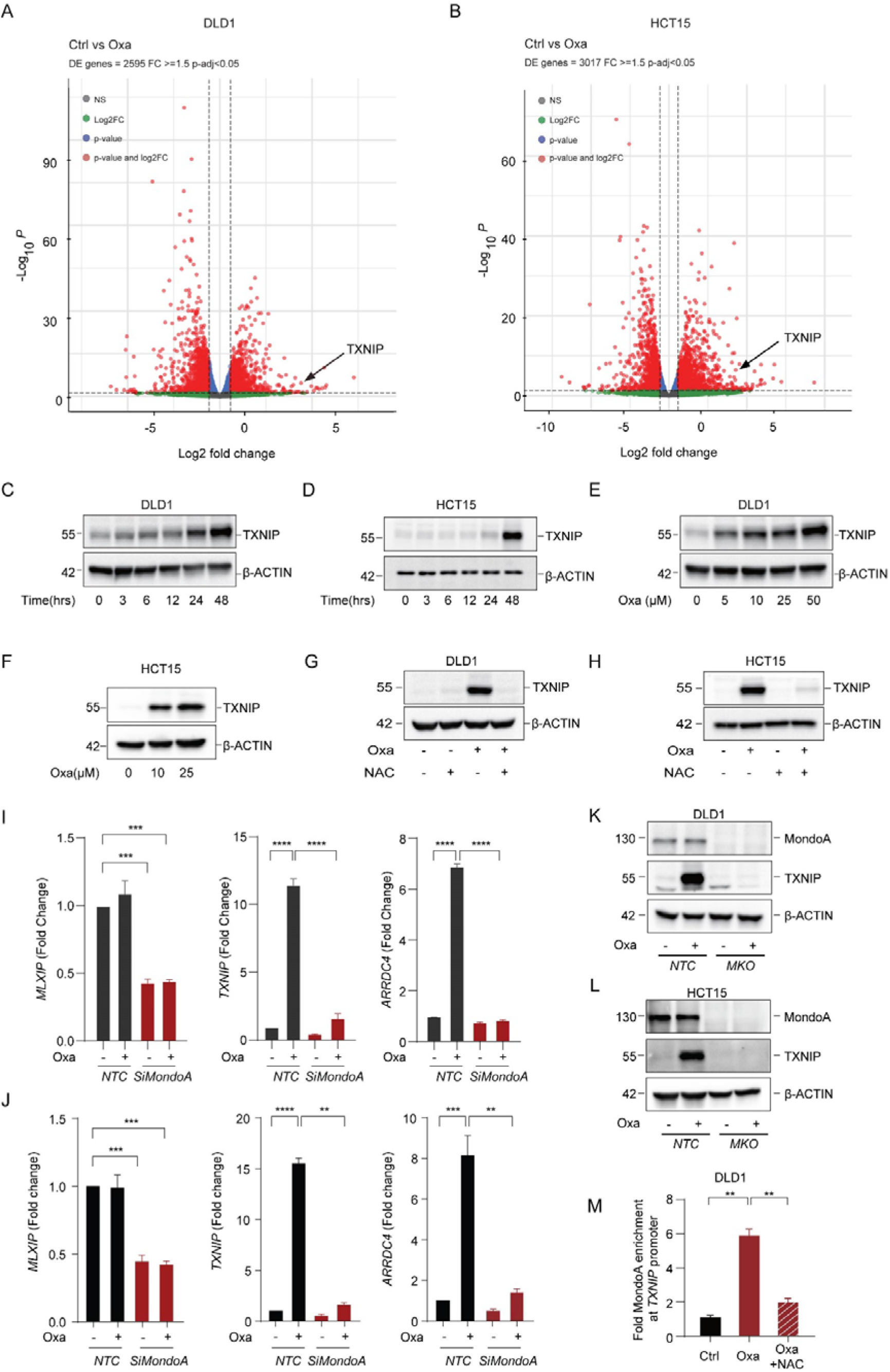
ROS mediates chemotherapy-induced TXNIP expression by modulating MondoA. (A-B) DLD1 cells (A) or HCT15 cells (B) were treated with 10 µM oxaliplatin for 48h and surviving cells were analysed by RNA sequencing. A volcano plot (log2 FC versus negative log of P value) was used to visualize statistically significant gene expression changes (fold ≥1.5 and adjusted P value <0.05). *TXNIP* is labelled. The number of DE genes is indicated in the upper left. 3 biological replicates per group. (C-D) Western blotting analysis of TXNIP expression in DLD1 cells (C) or HCT15 cells (D) treated with oxaliplatin at different time points. β-ACTIN was used as an internal reference. (E-F) Western blotting analysis of TXNIP expression in DLD1 cells (E) or HCT15 cells (F) treated with oxaliplatin at different doses for 48h. (G-H) Immunoblot analysis of TXNIP in DLD1 cells (G) or HCT15 cells (H) treated with N-acetyl-L-cysteine (1.25mM) or oxaliplatin (10µm) or the combinational treatment for 48h. (I-J) Quantification of MLXIP (MondoA), TXNIP and ARRDC4 mRNA in DLD1 cells (I) or HCT15 cells (J) upon knockdown of MLXIP by siRNA after treatment with 10µm oxaliplatin treatment for 48h. (K-L) Immunoblot analysis of TXNIP expression in MondoA-knockout DLD1 cells (K) or HCT15 cells (L) after 10µm oxaliplatin treatment for 48h. (M) MondoA occupancy on the promoters of *TXNIP* in DLD1 cells treated with 10µm oxaliplatin or the combinational treatment with NAC (1.25mM) for 48h. Results shown, excluding A and B, are representative of three independent experiments. All values were expressed as mean ± SEM. Two-tailed Student’s t test; **p<0.01, ***p < 0.001, ****p < 0.0001, vs. Control.

The thioredoxin (Trx) antioxidant system includes NAPDH, thioredoxin reductase (TrxR), and Trx. TXNIP is essential for redox homeostasis due to its ability to bind to Trx and inhibit Trx function and expression^47^. As discussed, oxaliplatin treatment induces ROS^48^, whilst oxidative stress is associated with TXNIP expression^49^. As such, we considered whether the increase in TXNIP expression after oxaliplatin treatment was mediated by ROS. In line with previous studies^50^, oxaliplatin was observed to increase ROS production in DLD1 and HCT15 cells (Figure S3A-B). Next, to understand whether it was this increase in oxaliplatin-dependent ROS that drove the increase in TXNIP, we administrated N-acetyl-L-cysteine (NAC, reactive oxygen species inhibitor) with oxaliplatin and observed no increase in TXNIP expression (Figure 2G-H, S3C).

We next investigated which transcription factor may mediate ROS-induced TXNIP expression. The RNA-seq data revealed 23 differentially expressed genes (DEGs) shared between both cell lines, including TXNIP (Figure S3D-E, Suppl Table 2). One of these DEGs, was arrestin domain-containing protein 4 (*ARRDC4*). ARRDC4 was increased after oxaliplatin treatment (Figure S3D-E; validated by RTPCR [Figure S3F-G]), and, like TXNIP, this increase shown to be dependent on ROS (Figure S3F-G). TXNIP and ARRDC4 are paralogs showing 63% similarity and are both regulated by the transcription factor MondoA^51,52^, indeed TXNIP and ARRDC4 have been reported to be highly MondoA-dependent^53^. We therefore assessed MondoA expression before and after oxaliplatin treatment, finding no change (Figure S3H). With MondoA having previously being shown to shuttle into the nucleus to carry out its functions^53^, we assessed for MondoA in different cellular fractions. The result showed MondoA was indeed translocated into the nucleus after oxaliplatin treatment (Figure S3I).

To assess the role of MondoA in oxaliplatin-induced TXNIP upregulation, we established MondoA-KO cells and MondoA-KD cells using CRISPR-Cas9 and siRNA, respectively. Using these models we saw that the removal or decrease of MondoA resulted in the loss of increased expression of both TXNIP and ARRDC4 after oxaliplatin treatment (Figure 2I-L). To further strengthen our conclusions, we used ChIP-PCR to verify the dependence of these processes on MondoA. Relative to the control, the amount of MondoA on the TXNIP promoter was significantly increased after oxaliplatin treatment, which was compromised after combined treatment with NAC (Figure 2M). Taken together, these results demonstrated that ROS production was responsible for oxaliplatin-induced TXNIP overexpression by activating MondoA transcriptional activity.

### TXNIP regulates the expression and secretion of GDF15

TXNIP has been reported to regulate both the innate and adaptive arms of the immune system^54^. In support of this, we found TXNIP expression to be positively associated with the expression of T cell markers, antigen presentation and cytokine transcripts when using the COAD TCGA dataset (Figure S4A). The enrichment of TXNIP in the cytoplasm indicated that TXNIP may mediate anti-tumor effects by regulating immunologically relevant cytoplasmic processes (Figure S4B-C)^55^, for example, the NLRP3 inflammasome^55^. The formation and activation of the NLRP3 inflammasome leads to self-cleavage and activation of caspase 1, which in turn promotes the release of the pro-inflammatory cytokine IL-1β. However, the correlation between TXNIP and IL-1β or IL-18 was not significant (Figure S4A). Similarly, knockout of TXNIP led to no alteration in caspase 1 activation and IL-1β production (Figure S4D-E), with no detectable IL-1β protein in the supernatants, suggesting TXNIP failed to activate the NLRP3 inflammasome upon chemotherapeutic treatment.

We therefore considered whether TXNIP may be capable of regulating the expression and/or secretion of other immunologically-relevant soluble factor(s) from the epithelial cell. To this end, we performed mass-spectrometric analysis of supernatants collected from non-targeting control (NTC) and TXNIP-KO (TKO) DLD1 cells and identified a total of 832 proteins from the conditional media and 157 differentially expressed soluble proteins (p<0.05). Protein data can be found from Supplementary Table 3. Growth/Differentiation Factor 15 (GDF15) was the most highly differentiated secreted protein associated with TXNIP loss (Figure 3A). This result was confirmed using a cytokine array, where GDF15 was additionally seen to be secreted at lower levels in response to oxaliplatin; in line with the upregulation of TXNIP (Figure 3B). These results showed that oxaliplatin decreases GDF15 secretion in a TXNIP dependent manner, and that the knockout of TXNIP alone could drive the secretion of GDF15. Intriguingly, other factors were seen to be altered in a similar manner, for example plasminogen activating inhibitor (PAI-1; SERPINE1. Figure 3B Row I, columns 1 and 2) suggestive of a TXNIP dependent signature which is broadly indicative of wound-healing, however with the proteomics showing GDF15 as the dominant factor, we focussed on this pathway.

**Figure 3.**
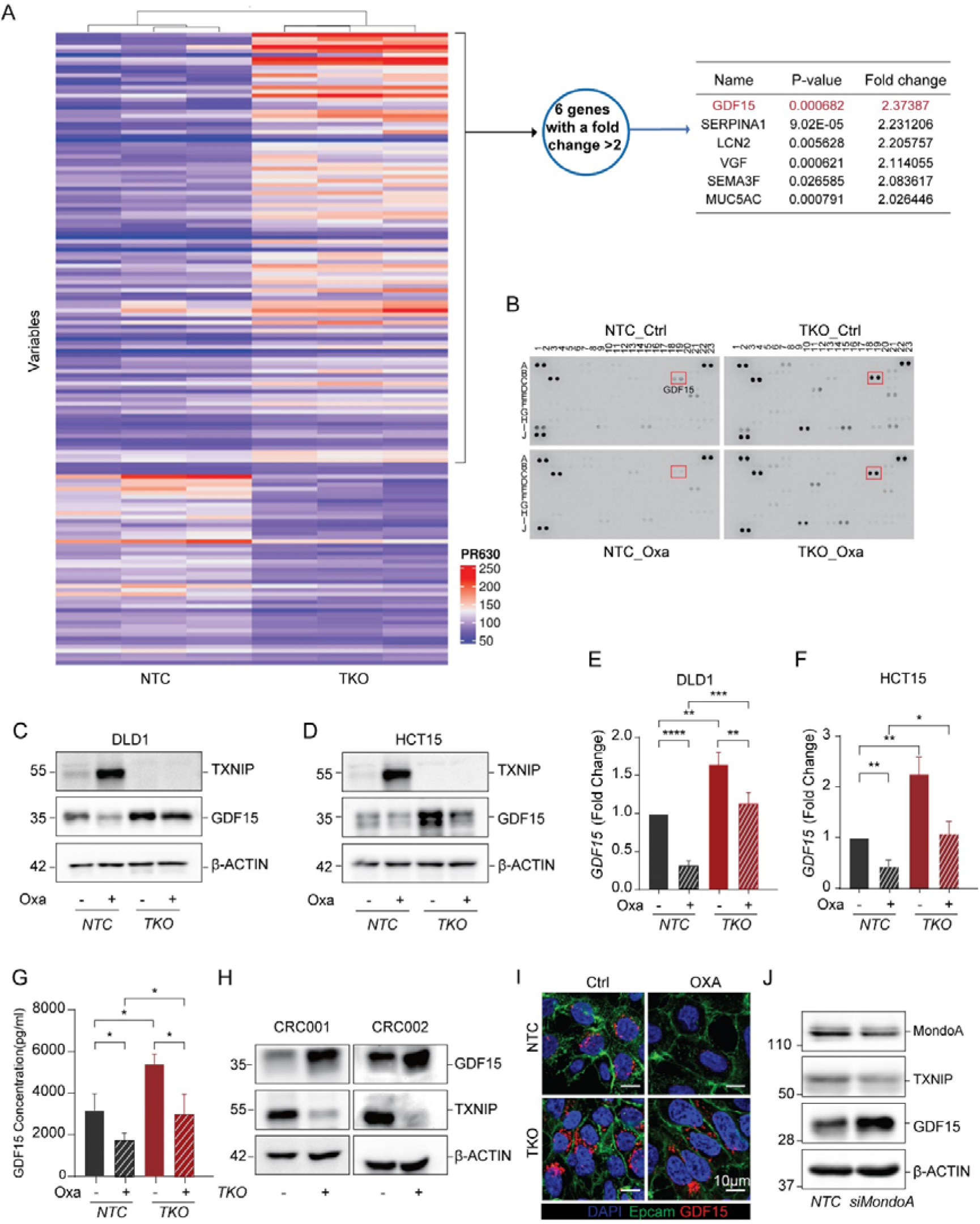
TXNIP regulates GDF15 expression. (A) Proteomic analysis of the conditional media from TXNIP-KO (TKO) and control (NTC) DLD1 cells as assessed by mass spectrometry. Heatmap illustrating differentially expressed proteins (left panel) and table showing the top six upregulated proteins in conditional media from TKO cells (right table). (B) 105 plex cytokine arrays incubated with conditional media from TKO and NTC cells with or without 10µM oxaliplatin treatment for 48h. The respective GDF15 spot is highlighted (red box). (C-D) Immunoblotting of TXNIP and GDF15 in NTC and TKO DLD1 cells (C) and NTC and TKO HCT15 cells (D) with or without drug treatment (10µm oxaliplatin for 48h). (E-F) Pooled densiometry data from 3 repeats of C and D. Standard error bars shown. (G) GDF15 concentration in conditional media for E were determined by ELISA. Standard error bars are shown. (H) Immunoblot of TXNIP and GDF15 in NTC (TKO-) and TKO (TKO+) PDTOs: CRC001 (left panel), CRC002 (right panel). (I) Immunofluorescent detection of GDF15 in NTC and TKO DLD1 cells with or without 10µm oxaliplatin treatment for 48h as assessed by confocal microscopy. DAPI (blue), Epcam (green), GDF15 (red). (J) Immunoblotting of MondoA, TXNIP and GDF15 in MondoA-knockdown (siMondaA) and control (NTC) DLD1 cells. Results shown are representative of three independent experiments. All values were expressed as mean ± SEM. Two-tailed Student’s t test; *p<0.05, **p<0.01, ***p < 0.001, ****p < 0.0001, vs. Control.

Having established the dependence of GDF15 on TXNIP, we next assessed the effects of oxaliplatin treatment on GDF15. The downregulation of GDF15 was more pronounced at later time points and higher drug dosages; the opposite trend to TXNIP (Figure S5A-B). Using Western blotting we showed that TXNIP knockout rescued the inhibitory effects of oxaliplatin on GDF15 expression in DLD1 cells (Figure 3C, E1), with a similar pattern being observed in TXNIP-KO HCT15 cells (Figure 3D, F). In contrast, TXNIP-overexpressing DLD1 cells showed lower GDF15 expression compared to control cells (Figure S5C-D). We quantitated soluble GDF15 concentrations by ELISA finding >5ng/ml in the supernatant of TXNIP-KO cells (Figure 3G), whilst a higher expression of GDF15 was also detected in TXNIP-KO PDTOs (Figure 3H). Next, using confocal imaging, we observed GDF15 was enriched in the cytoplasm in untreated cells, suggestive of it being stored in secretory granules, with no staining seen after oxaliplatin treatment. In line with immunoblot analysis, confocal imaging showed TXNIP-KO cells expressed more GDF15, which, unlike the control, was retained after oxaliplatin treatment (Figure 3I, S5E).

As ROS mediated the activation of the MondoA-TXNIP axis, we aimed to assess the effect of these factors on GDF15 expression. In line with our previous findings, knocking down MondoA decreased the expression of TXNIP, but increased GDF15 expression (Figure 3J, S5F), suggesting the involvement of MondoA in the regulation of GDF15 expression. Furthermore, pre-incubation of the target cells with NAC abolished the suppression of GDF15 by oxaliplatin, which was partially rescued by overexpressing TXNIP (Figure S5F), suggestive of the important role of ROS in GDF15 regulation. Collectively, these data demonstrated the activation of MondoA by ROS modulates both TXNIP and GDF15.

### GDF15 expression is upregulated in CRC and associated with poor prognosis

Consistent with previous reports^34^, GDF15 was observed to be upregulated in CRC tumor samples in comparison with normal tissue or epithelial cells by both TCGA COAD and scRNA epithelial sequencing analyses respectively (Figure S6A-B). This observation was validated by IHC staining (Figure 4A-B). To assess if the inverse relationship of TXNIP and GDF15 we had observed *in vitro* could be observed *in situ*, we assessed relative transcriptomic and protein expression using the TCGA COAD dataset and historic patient samples respectively, finding the same inter-relationship (Figure 4C-D). Using the same pre-T and post-T fresh patient samples described in Figure 1, we probed for GDF15, finding decreased GDF15 expression after treatment, except for the same three aggressive cases which had previously been shown to show no increased TXNIP expression (Figure 4E-G).

**Figure 4.**
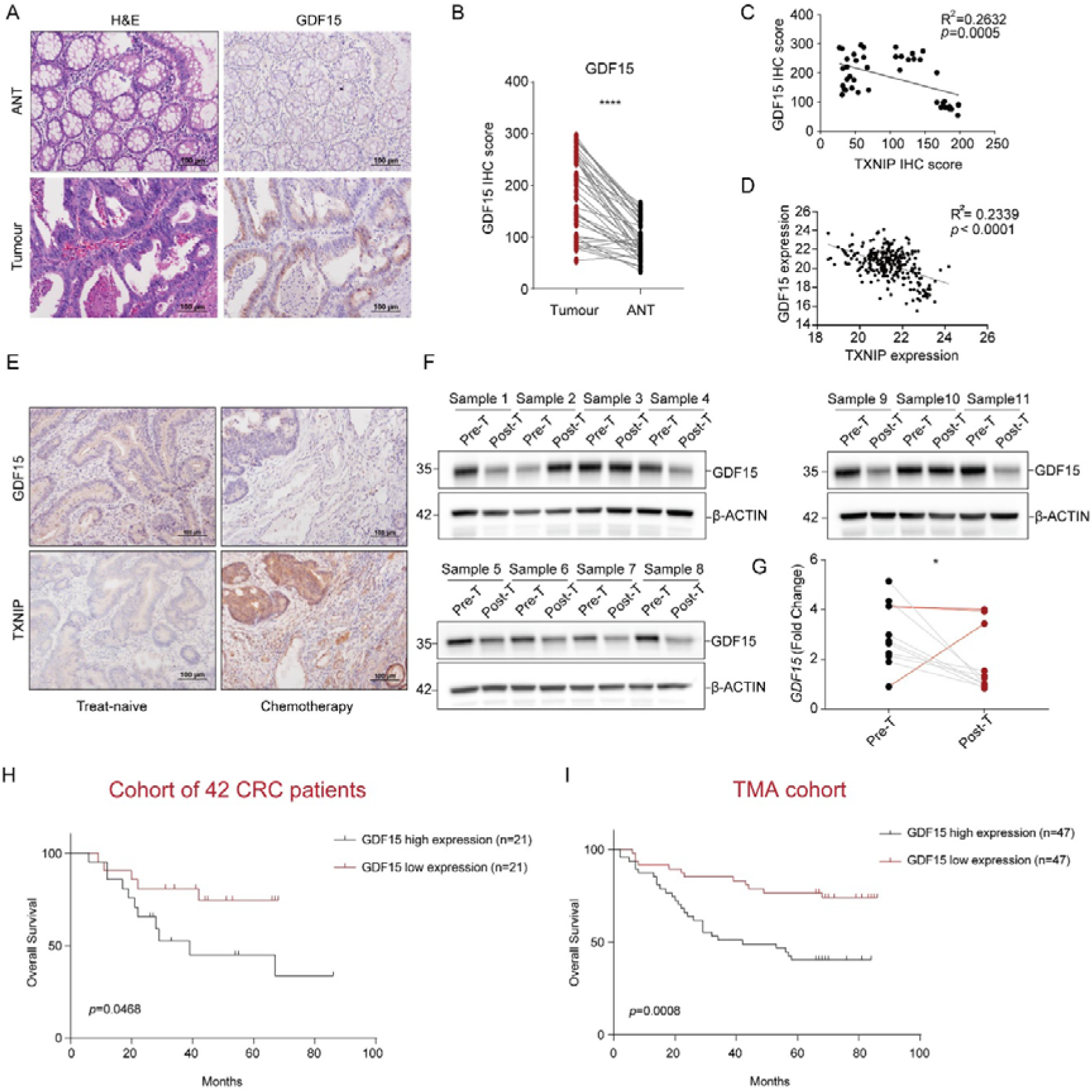
Higher GDF15 expression is observed in CRC tumor samples, however it is decreased post-chemotherapeutic treatment. High levels of GDF15 are associated with poor prognosis. (A) Detection of GDF15 in both tumor and adjacent normal tissue (ANT) samples from patients with primary colorectal cancer. Magnification ×200. (B) Statistical analysis of GDF15 IHC score between ANT and tumor tissue (n=42). (C-D) Correlations of TXNIP and GDF15 protein(cohort of 42 CRC patients) (C) and TXNIP and GDF15 transcripts (TCGA COAD) (D). Pearson correlation coefficients (R^2^) are indicated. (E) Sequential sections from colorectal tumor samples collected pre- and post-neo-adjuvant chemotherapy. Detection of TXNIP and GDF15 by IHC. (F) GDF15 expression in 11 paired treatment-naïve (Pre-T) tumor samples and oxaliplatin-based neo-adjuvant chemotherapy treated tumor samples (Post-T). (G) GDF15 mRNA levels in samples from F (aggressive cases highlighted in red). (H-I) Kaplan–Meier analysis of overall survival in CRC patients with different GDF15 staining scores from a cohort of 42 CRC patients (H) and CRC tumor tissue microarray (n=94) (I). Results shown are representative of three independent experiments. All values were expressed as mean ± SEM. *p<0.05, ****p < 0.0001, vs. Control.

We then sought to understand the clinical relevance of GDF15 in CRC. When assessing for the impact of increased expression of GDF15 on survival, we found associations between low GDF15 and improved outcome at the protein level in tissue (Figure 4H-I), and in two independent public transcriptomic datasets (Figure S6C,D), suggesting that GDF15 contributes to tumor progression in CRC. In an opposite manner to TXNIP, GDF15 showed a significantly positive correlation with clinical stage and lymph node metastasis in CRC specimens (Table 3 and Table 4)

**Table 3:** Association between GDF15 expression and clinicopathological features of patients with colorectal cancer in the cohort of 42 CRC patients.

|  | Total<br>(n = 42) | GDF15 expression |  | *P* value |
| --- | --- | --- | --- | --- |
|  |  | Low<br>(n = 21) | High<br>(n = 21) |  |
| Gender |  |  |  | >0.9999 |
| Male | 22 | 11 | 11 |  |
| Female | 20 | 10 | 10 |  |
| Age(year) |  |  |  | 0.7171 |
| <65 | 32 | 15 | 17 |  |
| ≥65 | 10 | 6 | 4 |  |
| T stage |  |  |  | 0.4520 |
| T1-T2 | 9 | 6 | 3 |  |
| T3-T4 | 33 | 15 | 18 |  |
| N stage |  |  |  | 0.0219* |
| N0 | 14 | 11 | 3 |  |
| N1+N2+N3 | 28 | 10 | 18 |  |
| M stage |  |  |  | 0.0017* |
| M0 | 25 | 18 | 7 |  |
| M1 | 17 | 3 | 14 |  |
| Clinical stage |  |  |  | 0.0219* |
| I/II | 14 | 11 | 3 |  |
| III/IV | 28 | 10 | 18 |  |

\**P*<0.05

**Table 4:**
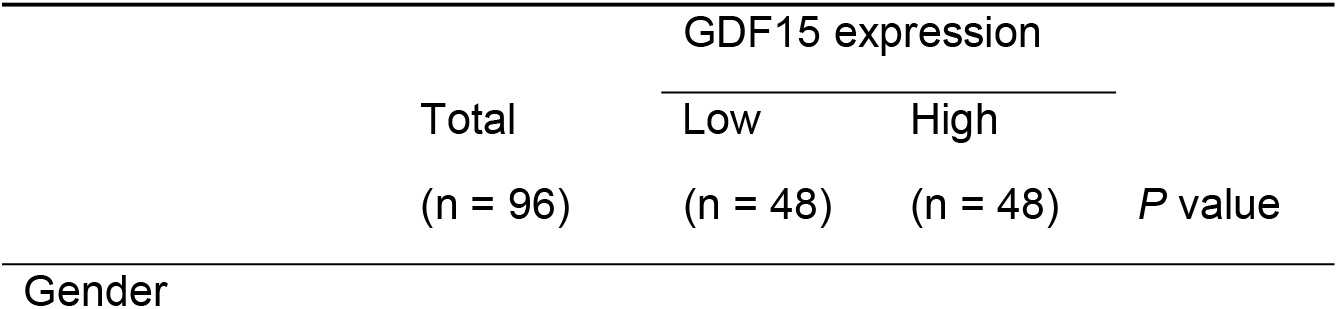
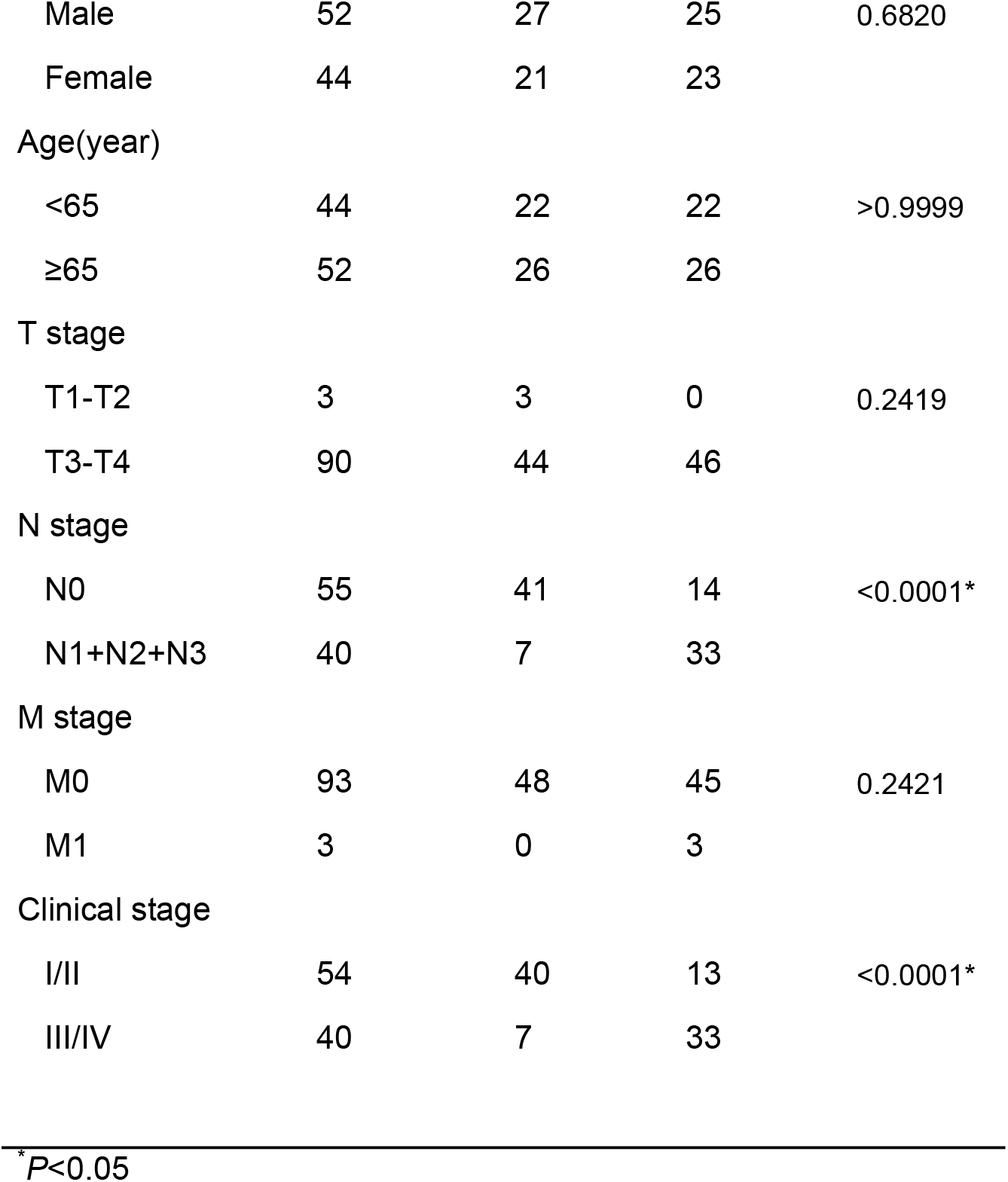
Association between GDF15 expression and clinicopathological features of patients with colorectal cancer in TMA cohort.

### The role of TXNIP-GDF15 axis in immune regulation

GDF15 has been reported to have multiple immunological impacts however some reports have been queried owing to the discovery of contaminating TGF-β1 in recombinant GDF15 preparations^56^. As such, to explore the immune impacts of GDF15, we opted to predominantly use cellular systems and resultant conditioned supernatant (Figure S7A-B).

When stimulating PBMCs with anti-CD3 and anti-CD28 in the presence of GDF15-enriched conditioned media from the TXNIP KO cell line, we observed a small but significant decrease in cell number, that was reversed using supernatant from a TXNIP / GDF15 double KO cell line (Figure 5A-B). Further analysis showed that both CD8 and CD4 T cell proliferation was inhibited by GDF15 enriched supernatant (Figures 5C-F), with IFNγ concentrations in the supernatant also being seen to lower (Figure 5G-H).

**Figure 5.**
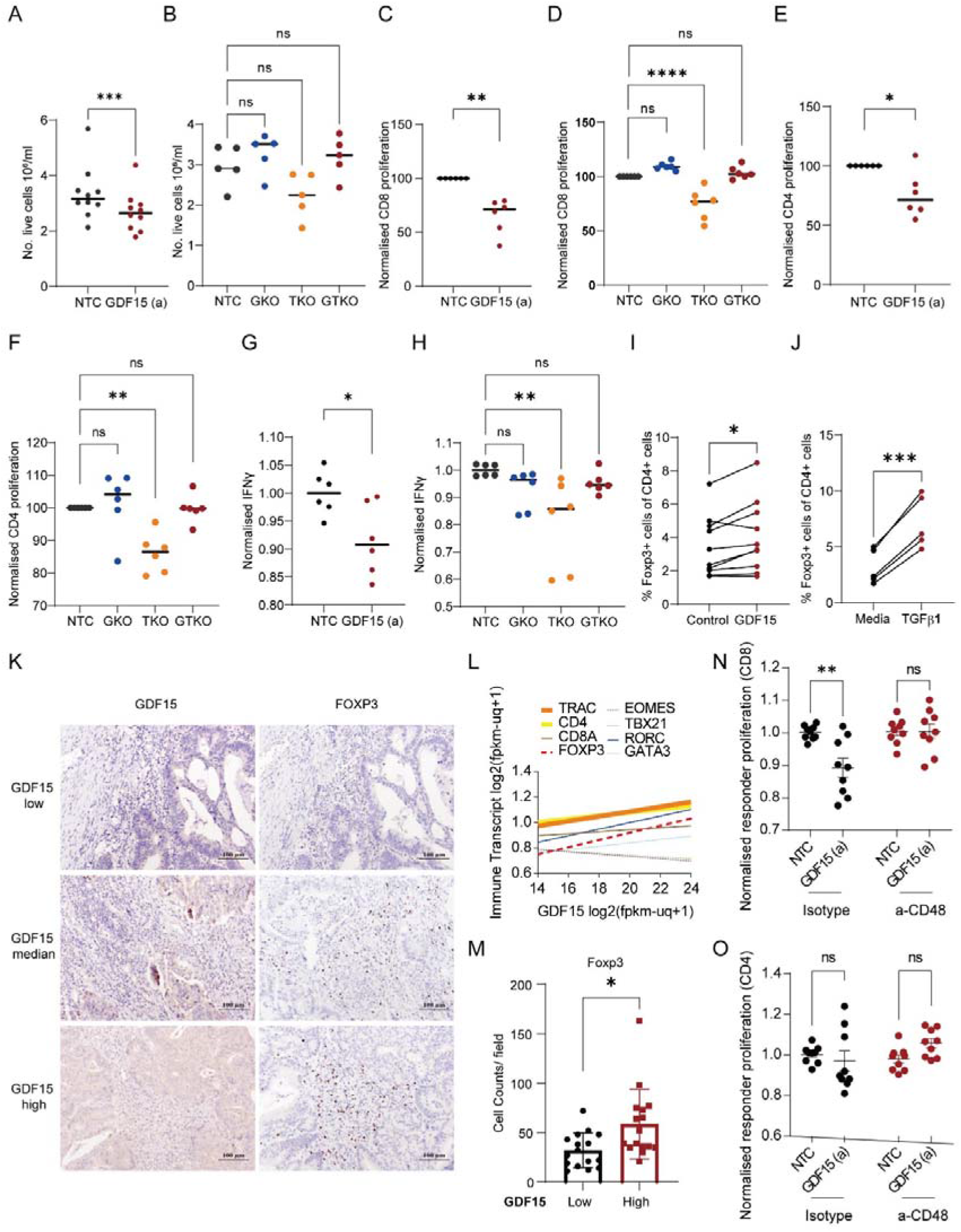
GDF15 induces Tregs in a CD48 dependent manner. (A-B) PBMCs were stimulated with anti-CD3 and anti-CD28 for 4 days in the presence of fresh supernatant from indicated cell lines (NTC,GKO,TKO,GTKO; GDF15a). Live cells were counted using trypan blue and a haemocytometer. n=10 (A) and n=5 (B). (C-F) Labelled PBMCs were stimulated with anti-CD3 and anti-CD28 for 4 days in the presence of fresh supernatant from indicated cell lines, before being stained with anti-CD3 and anti-CD8 (C-D) or anti-CD4 (E-F) antibodies and measured by flow cytometry. Normalised proliferation on gated CD3^+^CD8^+^ or CD3^+^CD4^+^ cells is shown. n=6. (G-H) Normalised IFNγ concentrations in the supernatant of cells from C-F. (I-J) PBMCs were stimulated with anti-CD3 and anti-CD28 for 4 days in the presence of fresh supernatant from NTC or GDF15a cell lines (I) or media alone or 5ng/ml recombinant human TGFβ1 (J). Cells were stained with anti-CD3, anti-CD4 antibodies extracellularly before intranuclear staining of Foxp3 was performed. % of CD4^+^Foxp3^+^ cells are shown. n=10 (I) and n=5 (J). (K) Immunohistochemistry using anti-GDF15 and anti-Foxp3 antibodies on serial sections from colorectal cancer cases. (L) Correlations of indicated immune transcripts (normalised for PTPRC[CD45] expression) and GDF15 transcripts from TCGA COAD dataset. Thick line indicates R^2^ value >0.1 and dashed line indicates transcription factor. (M) Pooled data from K showing Foxp3^+^ cell counts in GDF15^low^ and GDF15^high^ populations; median split. n=32. (N-O) Isolated naïve CD4 cells were stimulated with anti-CD3 and anti-CD28 for 4 days in the presence of indicated cell line supernatant and either isotype control (10µg/ml) or anti-CD48 (10µg/ml) as indicated. These cells were then co-cultured with anti-CD3 stimulated proliferation dye labelled responder PBMCs for 4 days, before cells were stained for CD3, CD8 and CD4. Normalised proliferation dye (MFI) of the indicated responder population is shown. n=9. All values were expressed as mean ± SEM. *p < 0.05, **p < 0.01, ***p < 0.001, ****p < 0.0001, vs. Control.

A recent paper has shown that GDF15 is able to drive the differentiation of regulatory T cells (Tregs) from naïve CD4s via CD48 ligation^42^. Working on the hypothesis that it was Tregs that were inhibiting the T cell proliferation and IFNγ release within the mixed PBMC population, we observed a GDF15-dependent increase of Foxp3 within the CD4 pool (Figure 5I), however to a much lesser extent than when using active TGFβ1 (Figure 5J). To support these data we assessed for associations between GDF15 and FOXP3/Foxp3 in TCGA COAD dataset and our historic 42 patient cohort respectively, finding a significantly positive correlation between GDF15 and FOXP3 and enrichment of Foxp3 in the GDF15 high cases (Figure 5K-L). Finally, when stimulating naïve CD4 T cells in the presence of GDF15 enriched supernatant we were able to both differentiate these cells into functional Tregs and also block this functionality using an anti-CD48 antibody (Figure 5M-N).

### Loss of TXNIP/GDF15 axis functionality in advanced disease and the use of pre-treatment GDF15/TXNIP ratio as a biomarker of clinical response

With high GDF15, Treg infiltration and CD8 T cell dysfunction all being shown to be associated with poor prognosis in CRC^57–59^, and with the vast majority of CRC patients being treated with oxaliplatin, we next considered whether the TXNIP/GDF15 axis, an axis which should regulate these processes to the benefit of the patient, remained functional in metastatic disease. In the course of this project we had observed a clear distinction in the TXNIP/GDF15 response to oxaliplatin when looking at cell lines derived from primary and secondary sites (Figure 6A-B). This difference can be seen most clearly when assessing the ratio of GDF15 to TXNIP (GDF15/TXNIP) pre-treatment (Figure 6C). We next assessed if there was a difference in the correlation between TXNIP and GDF15 in metastatic and primary disease, finding the significant inverse relationship in primaries discussed earlier was lost in metastatic samples (Figure 6D-E). As resistance to chemotherapy is commonly observed in patients with metastatic disease, we developed two oxaliplatin-resistant lines, finding that they also lost oxaliplatin-induced TXNIP/GDF15 responsiveness (Figure 6F-G), with GDF15/TXNIP ratios strongly resembling those of the cell lines derived from different sites (Figure 6H).

**Figure 6.**
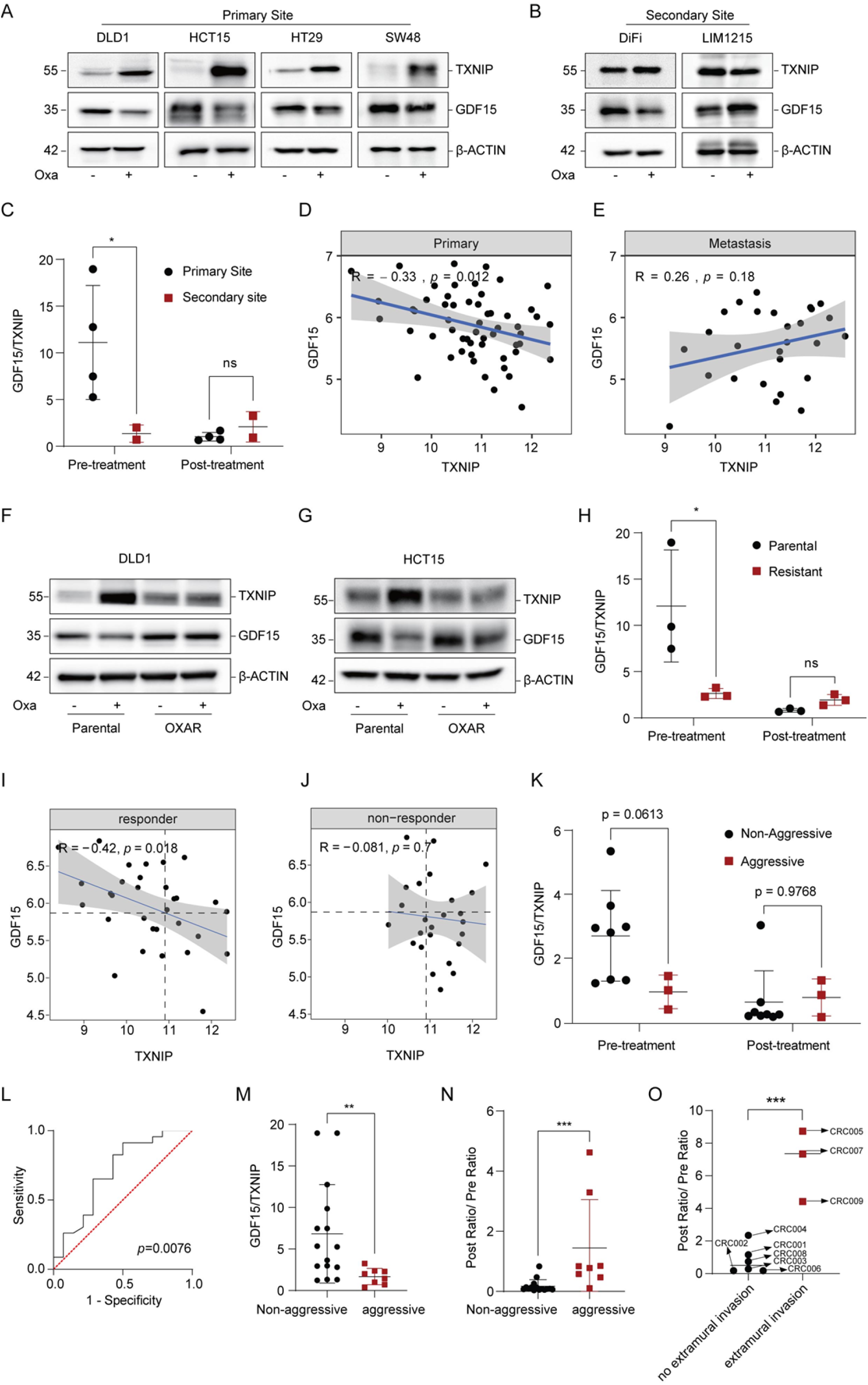
Loss of a oxaliplatin responsive TXNIP/GDF15 axis is associated with advanced disease and chemotherapeutic resistance, and pre-treatment GDF15/TXNIP ratio can be used as a biomarker of treatment response. (A-B) Immunoblot analysis of TXNIP and GDF15 expression after 48h of 10µm oxaliplatin treatment in colorectal cancer cell lines, including DLD1, HCT15, HT29, SW48 (A, derived from primary site), and DiFi, LIM1215 (B, derived from secondary site). (C) Ratio of GDF15/TXNIP for cell lines in A-B treated as indicated and measured using densiometry. (D-E) Microarray data showing the correlation between GDF15 and TXNIP mRNA expression in primary (D) or metastatic (E) CRC tumors. R and p values shown (Pearson’s). (F-G) Immunoblot analysis of TXNIP and GDF15 expression after 48h of 10µm oxaliplatin treatment in oxaliplatin-resistant (OXAR) cells: DLD1-OXAR (F) and HCT15-OXAR (G). (H) Ratio of GDF15/TXNIP for cell lines in F-G treated as indicated and measured using densiometry. (I-J) Microarray data showing the correlation between GDF15 and TXNIP mRNA expression in primary tumors that respond (responder; I) or do not respond (non-responder; J) to FOLFOX chemotherapy. R and p values shown (Pearson’s). (K) Ratio of GDF15/TXNIP for primary tumours in Figures 1D and 4F treated as indicated as measured using densiometry. (L) Receiver operating characteristic (ROC) curve showing area under the curve and p values for the use of pre-treatment GDF15/TXNIP ratio in predicting responsiveness to oxaliplatin (O; responder [n=23] and non-responder [n=14]) using publicly available data. (M) Pooled pre-treatment data (ratio of GDF15/TXNIP) from C, H, K with ‘aggressive’ classed as secondary site, resistant to oxaliplatin and aggressive and ‘non-aggressive’ primary site, sensitive to oxaliplatin and non-aggressive (N) Post-treatment GDF15/TXNIP ratio divided by pre-treatment GDF15/TXNIP ratio for C, H, K. ‘Aggressive’ and ‘Non-aggressive’ defined as in M. (O) Post-treatment GDF15/TXNIP ratio divided by pre-treatment GDF15/TXNIP ratio for patient derived organoids grouped into primary tumours with and without extra-mural invasion. * p<0.05 using Sidak’s multiple comparisons test (C, H, K) ** p<0.01 *** p<0.001 using Mann Whitney (M, N) or unpaired t test (O). Western results shown are representative of three independent experiments.

We next considered whether this oxaliplatin resistance-associated loss of TXNIP/GDF15 responsiveness could be observed in progressive primary tumors. We first assessed TXNIP-GDF15 correlations in primary samples where chemotherapeutic response was known (non-responder vs responder) finding the inverse ‘functional’ relationship was only present in responders (Figure 6I-J). We then assessed our pre-treatment and post-treatment fresh tumour samples finding similar ratios to those observed in the cell line models when splitting the cohort into aggressive and non-aggressive disease (Figure 6K). These data collectively suggest that the loss of the responsive TXNIP/GDF15 axis (oxaliplatin inducing ROS, driving TXNIP upregulation via MondoA, leading to a decrease in GDF15 secretion) is associated with both disease progression and chemotherapeutic resistance.

We then questioned whether or not the pre-treatment ratio of GDF15/TXNIP could be used as a potential biomarker of oxaliplatin treatment responsiveness. To test this hypothesis we first assessed whether or not the ratio could be used to differentiate cell lines from primary or secondary sites (Figure S8E) or oxaliplatin resistant lines from non-resistant (Figure S8F) or aggressive from non-aggressive tumours (Figure S8G) as controls. We then tested this ratio using a publicly available dataset finding that pre-oxaliplatin treatment GDF15/TXNIP ratio could be used to determine treatment response (Figure 6L). Interestingly this result was completely negated if oxaliplatin was combined with radiotherapy.

Finally, as the data clearly showed a differential in ratio change between pre and post treated ‘aggressive’ and ‘non-aggressive’ groups (definitions in the appropriate legend), we tested a new metric, post-treatment GDF15/TXNIP ratio divided by pre-treatment GDF15/TXNIP ratio (Figure S8H-J), to see if this would improve the overall differential. We found that by adopting the new metric not only did the combined differential increase (Mean of 6.82 vs 1.68 [fold change of 4.1] for single pre-treatment GDF15/TXNIP ratio against 0.05 vs 1.44 [fold change of 28.8] for the combined) but so did the significance (Figure 6M,N). Given that there are no publicly available datasets pre and post oxaliplatin treatment, we used organoids derived from primary tumours to test this new metric by measuring GDF15 and TXNIP pre and post treatment. Splitting the organoid groups into those with extra-mural invasion (considered more aggressive) and those without (less aggressive), we could see a significant difference between the groups (Figure 6O).

## Discussion

Colorectal cancer is the third most common cancer worldwide, with 1.9 million cases reported in 2020. Five year survival ranges greatly, from 13-88%, depending on stage at presentation, age and sex^60^. Chemotherapy, predominantly oxaliplatin-based, is the most common first line therapy and has been increasingly shown to be capable of turning a ‘cold tumor’ with low active immune infiltrate into a ‘hot tumor’ with improved infiltration. This conversion lays the foundation for current combinational chemo-immunotherapies, however, beyond innate stimulation through disease associated molecular patterns (DAMPs) and the presentation of neoantigens, our understanding into exactly how the immune system, especially the adaptive arm, is ‘reawakened’ is limited.

Although tumor suppressor genes (TSGs) are well known to function by targeting oncoproteins for degradation or inducing cell death per se., we have sought to understand the role of one particular TSG, TXNIP, in mediating chemotherapy-induced immunogenicity. Our interest in TXNIP stemmed from its reported role in regulating epithelial oxidative stress and its increased expression in fresh tumor samples after oxaliplatin treatment. By taking this observation, interrogating it *in vitro*, and investigating TXNIP’s role in regulating the TME, these data have revealed a previously unreported epithelial-immune axis, namely ROS-MondoA-TXNIP-GDF15-Treg. (Figure 7. Schematic diagram).

**Figure 7.**
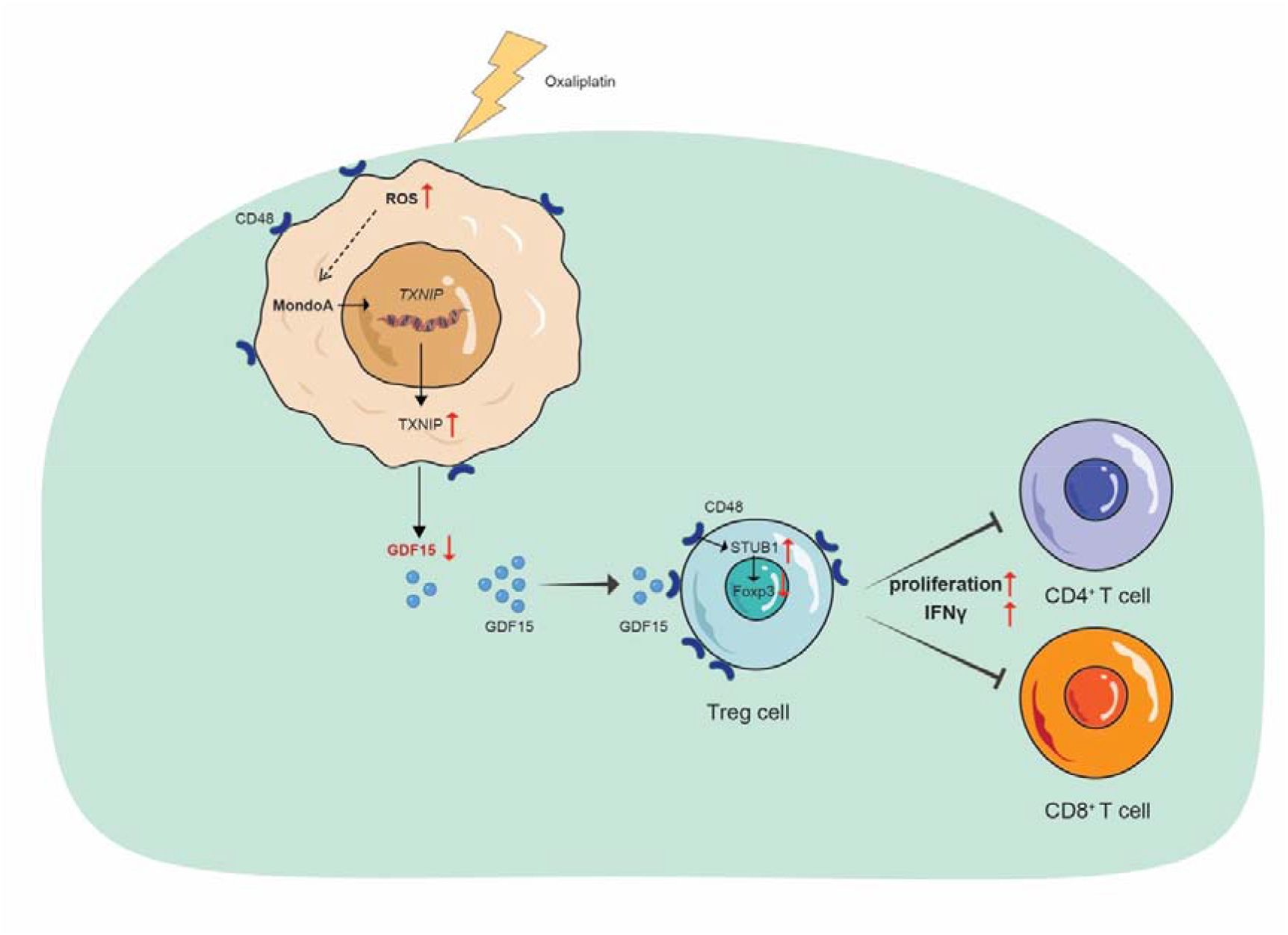
Schematic diagram of the underlying mechanism of oxaliplatin-induced immunogenicity by regulating MondoA/TXNIP/GDF15 signalling pathway in CRC.

The balance of reductive and oxidative processes is crucial for cellular life. Dysregulation can promote oxidative stress which contributes to diverse pathologies, including neurodegenerative disorders, autoimmune diseases and cancers. Intracellular ROS in tumor cells has been observed to increase upon chemo- and radiotherapy, leading to apoptosis^61^. Additionally, ROS levels in innate or adaptive immune cells are broadly associated with activation and anti-tumor effects^24,62,63^. A recent study by Gao *et al.* identified that the ROS induced by chemotherapy increased the secretion of HMGB1 to facilitate the infiltration of T cells^64^, highlighting the importance of ROS in mediating cancer-immune cross talk. In this study, we found oxaliplatin-induced ROS generation could activate MondoA which, in turn, induced TXNIP expression. Furthermore, combining mass spectrometry, proteomic array and genetically modified models (CRISPR-KO and CRISPR-activation), before verifying *in situ,* we revealed that the ROS/MondoA/TXNIP axis negatively regulated GDF15 expression and secretion.

GDF15 has previously been shown to promote ‘M2’ macrophage differentiation, inhibit NK cell function and dendritic cell maturation^65,66^, however, as described the purified recombinant tools used in these studies have been shown to be contaminated with active TGF-β1, raising concerns (as all these effects can be ascribed to this pleotropic cytokine)^56,67^. In this study, to avoid this issue, we prioritised the use of cellular systems for our immunological assays. A recent study, which used mass spectrometry to confirm the material they used was not contaminated with TGF-β1, found that recombinant GDF15 was able to induce and maintain Tregs via interaction with CD48 on naïve T cells^42^. Our findings support this concept, further adding tissue validation (the association of high GDF15 and FOXP3/Foxp3) and the potential of preventing this process using CD48 blockade. Given these data and the well-reported negative prognostic impacts of Tregs in tumors, including in CRC, and the positive impact of chemotherapy, we put forward the following model. 1. Chemotherapy either promotes cell death or induces oxidative stress and ROS formation in the cells that survive. 2. The cells that survive do so by increasing TXNIP expression to help alleviate the impact of chemotherapy-induced ROS (or naturally carry a high level of TXNIP and are selected for). 3. This high level of TXNIP inhibits GDF15 expression which consequently inhibits the local generation of Tregs from naïve CD4 cells. 4. This decrease in Tregs allows other T cells, especially CD8s, to function and help to eradicate the remaining tumor, facilitating a durable response.

One of the most intriguing aspects of this work is the impact of the post-chemotherapeutic change (TXNIP^low^GDF15^high^ to TXNIP^high^GDF15^low^), and the lack of change, on outcome. TXNIP is a known TSG and, as such, we show increased expression is associated with better prognosis, whilst the inverse is true for GDF15 (leading to the ongoing development of targeting drugs)^57^. These data suggest that the post-chemotherapeutic change, something validated in primary CRC cell lines, spheroids, PDTOs and, critically, patients themselves, is associated with positive outcome. The lack of responsiveness seen in cell lines derived from secondary sites, resistant models and fresh tumors taken from patients with more advanced disease, suggests that this axis is ‘broken’ in these contexts. These data are supported by publicly available transcriptomic data showing that the negative correlation, indicative of response, is not seen in either primaries that do not respond to chemotherapy or in metastases. As such, these collective data suggest that there is a subgroup of patients who intrinsically carry, or develop, a lack of responsiveness, raising the possibility of using biopsies as a stratification tool. Indeed we were able to demonstrate that the pre-treatment GDF15/TXNIP ratio was able to predict tumours that were responsive to oxaliplatin from those that were not.

Aware of the fact that the change in GDF15/TXNIP ratio pre and post treatment would likely give a better differential between aggressive and non-aggressive groups, and aware of the fact that pre and post treatment biopsies are often difficult to control and justify clinically, we combined these ratios and tested this new metric using organoids. Using this technique and this new parameter/metric (change in GDF15/TXNIP ratio pre and post treatment) we were able to demonstrate that organoids have potential as sentinels of oxaliplatin responsiveness and disease progression, With this knowledge it may well then be possible to predict oxaliplatin non-responders, using a single GDF15/TXNIP pre-treatment ratio (biopsy; transcript or protein), or a potentially more sensitive combined post-treatment / pre-treatment ratio (organoids; protein), and change treatment plans accordingly. Indeed this methodology is especially pertinent to the use of anti-GDF15 therapeutics, allowing their potential use early in disease. As such these data champion targeted, effective therapy through biological understanding and functional assessment.

## Materials and Methods

### Public dataset analysis

The cancer genome atlas (TCGA) was used to compare the differential expression of TXNIP/GDF15 between adjacent normal samples and cancer patient samples. Gene expression data from TCGA was downloaded from either the GDC data portal (https://www.genome.gov/Funded-Programs-Projects/Cancer-Genome-Atlas) or UCSC Xena functional genomics explorer (https://www.xenabrowser.net). Both colon adenocarcinoma (COAD) and rectal adenocarcinoma (READ) cohorts were included as colorectal cancer cases. Four public datasets were used in this study for prognostic analyses, including GSE29621, GSE38832, GSE6988, and GSE52735. These datasets were downloaded from the Gene Expression Omnibus (GEO, http://www.ncbi.nlm.nih.gov/geo). For the survival analysis, the continuous variables were dichotomized via the survminer R package, and the Kaplan-Meier curves were performed using the survival R package. To measure TXNIP and GDF15 expression in normal and tumor epithelial cells from paired samples at single-cell level, we used normalized scRNA-seq data from 10 paired samples from colorectal cancer patients deposited in GSE132465. Microarray data from responder and non-responder to FOLFOX therapy for primary and metastatic lesion was downloaded from GSE28702 and normalized using RMA and converted to the gene level using an appropriate average. ROC analysis for publicly available data was performed using rocplot.org^68^.

### Human samples

This study was approved by Peking university Third Hospital Medical Science Research Ethics committee (Reference number IRB00006761-M2022237) and was performed in accordance with the principle of the Helsinki Declaration II. Information of the human cohorts is provided in Supplementary Table 7 and 8. Two cohorts, including 42 CRC tissues with tumor tissue and corresponding adjacent normal tissues (Supplementary Table 7) and 11 CRC tissues with pre-and matched post-oxaliplatin-based chemotherapy (Supplementary Table 8), were retrospectively collected from May 2014 to March 2021.

A human colorectal cancer tissue microarray (TMA) purchased from Shanghai Outdo Biotech Company Ltd (Shanghai, China). All tissue samples were collected before chemotherapy treatment. The TMA contained 97 colorectal cancer samples and paired adjacent normal tissues collected from patients between 2009 and 2018 and were accompanied by patient clinical data. Patient information of TMA is provided in Supplementary Table 9.

### Immunohistochemical (IHC) staining

The tumor tissues excised during the operation were immediately placed in 10% formalin for fixation^69^. To begin with, FFPE slides were dewaxed and rehydrated. After antigen retrieval in 0.01 M sodium citrate buffer (PH 6.0) in a microwave for 20 min, slides were treated with peroxidase block for 5 min and protein block solution for another 5 min at RT. Then Slides were incubated with primary antibody against TXNIP (Abcam, ab188865; 1:250), GDF15 (Protein-tech, 27455-1-AP; 1:500) and FOXP3 (Abcam, ab215206; 1:1000) overnight at 4°C. Post primary antibody incubation, tissues were incubated with secondary antibodies (EnVision Chem Detection Kit, DaKo Cytomation) at room temperature for 30 min. Followed by incubation with horseradish enzyme-labelled streptavidin solution for 10 min and then stained with DAB and haematoxylin. The stained tissues were interpreted by two pathologists blinded to the clinical parameters. Staining percentage scores were defined as: expression intensity × expression area. Expression intensity was scored from 0 to 3 (10 × 20 magnification, 5 different random fields of view were selected), representing negative, weakly staining (light yellow), moderately staining (pale brown with light background), and strongly staining (dark brown without background), respectively. Expression area was scored from 0 to 4: 0 (1–5%), 2 (26–50%), 3 (51–75%) and 4 (>75%). representing <5, 6–25, 26–50, 51–75, and, respectively. The degree of positive staining: 1–3 was classified as weakly positive (+); 4–6 as moderately positive (++); and 7–12 as strongly positive (+++). The intraclass correlation coefficient (ICC) analysis was used for assessing the level of agreement between independent reviewers. The ICC scores were 0.893, 0.912 and 0.905 for samples stained with anti-TXNIP, anti-GDF15 and anti-FOXP3 antibodies, respectively.

### scRNA-seq analysis

For comparing GDF15 expression in colorectal cancer tumor samples, we used log transformed-normalized single-cell RNA sequencing data derived from 63 colorectal cancer patients^70^ deposited at the Synapse (syn26844071) and extracted only tumor cells.

### Western blot

Cells were seeded into 6-well plates (4 × 10^5^ cells per well). The following day cells were replaced with fresh media for 1 hour and then treated as indicated in the Figures. Cell fractionation was performed with NE-PER™ Nuclear and Cytoplasmic Extraction Reagent (Thermo Fisher scientific, 78833), buffers were added with protease and phosphatase inhibitors.Following two washes with PBS, cells were lysed in 150-200 µl 1.5×sample lysis buffer (Table 5, 5×sample lysis buffer diluted in ddH_2_O). Cell lysates were measured using the BCA assay (Pierce™ BCA Protein Assay Kit, 23227) and run on SDS–PAGE with 30ug protein loaded. After blocking in 5% milk or 5% bovine serum albumin (BSA) in tris-buffered saline and Tween-20 (TBST) for 2 h at room temperature. Antibodies against MondoA (Abcam, 1:1000), IL-1β (1:1000, Cell Signaling Technology), Caspase 1(1:1000, Cell Signaling Technology), TXNIP (1:1000, Cell Signaling Technology), Cas9 (1:1000, Santa Cruz), GDF15 (1:1000, Abcam), β-Actin (1:5000, Proteintech), GAPDH (1:5000, Proteintech) and Lamin A (1:1000, Cell Signaling Technology) were used for incubation overnight at 4 °C.

**Table 5.** 5× sample lysis buffer.

| Reagent | Volume |
| --- | --- |
| 1M Tris PH6.8 | 2.5ml |
| SDS | 1g |
| Glycerol | 5ml |
| Hit up to 60-70 degree |  |

### Cell lines and reagents

Human colon adenocarcinoma cell lines DLD1, DiFi, and SW48 were purchased from ATCC. LIM1215 was a generous gift from Dr. Sabine Teipar (University Leuven, Belgium). HT29 and HCT15 were generous gifts from Dr. Juan Jose Garcia Gomez (University College London). DLD1, HCT15, HT29, and LIM1215 were maintained at 37°C with 5% CO_2_ in RPMI supplemented with 10% fetal bovine serum (FBS), 1% Penicillin/streptomycin (P/S), and L-glutamine (2 mM). DIFI and SW48 were grown at 37°C with 5% CO_2_ in DMEM supplemented with 10% FBS, 1% P/S and L-glutamine (2 mM). All the CRC cell lines tested negative for mycoplasma throughout the study.

### RNA sequencing

The RNA-Seq experiments were performed by Novogene (Cambridge, UK) Company Limited^71^. Briefly, total RNA from CRC cells was isolated using TRIzol reagent. Messenger RNA was purified from total RNA using poly-T oligo-attached magnetic beads. After fragmentation, the first strand cDNA was synthesized using random hexamer primers followed by the second strand cDNA synthesis. The library was ready after end repair, A-tailing, adapter ligation, size selection, amplification, and purification. For the data analysis, base calls were performed using CASAVA. Reads were aligned to the genome using the split read aligner TopHat (v2.0.7) and Bowtie2, using default parameters. HTSeq was used to estimate abundance.

### Transfection

For transient transfection, siRNA was transfected into different cell lines using Lipofectamine™ RNAiMAX Transfection Reagent (Thermo Fisher Scientific, 13778075). 3 × 10^5^ cells were seeded in 6-well plates in antibiotic-free complete medium. After 24 h, 5 µl of Lipofectamine™ RNAiMAX Transfection Reagent and 25 pM siRNA (Table 6) were mixed thoroughly and incubated for 20 min before added to the cells at room temperature. Knockdown efficiency was assessed by western blot and PCR analysis after 48 h.

**Table 6:** Sequence of siRNA oligonucleotides.

| Reagent or Resource | Source | Identifier |
| --- | --- | --- |
| ON-TARGETplus non-targeting pool siRNA | Dharmacon | D-001810-10-05 |
| ON-TARGETplus SMARTpool siRNA Human MLXIP | Dharmacon | L-008976-00-0005 |

### RNA isolation and quantitative real-time PCR (qRT-PCR)

Cells were lysed in 0.7 ml of TRIzol Lysis Reagent (Invitrogen, 15596026), vortexed and incubated for 10 min at room temperature. RNA was extracted using the RNeasy Mini Kit (Qiagen, 74104) in the presence of RNase-free DNase (Qiagen, 79254). cDNA was synthesized by reverse transcription using a SuperScript™ II Reverse Transcriptase kit (Thermo Fisher scientific, 18064022). qRT-PCR was performed with Power SYBR green PCR master mix (Applied Biosystems, 4309155). Primers are listed in Table 7. Data analysis was conducted with the QuantStudio 6 Flex Real-Time PCR System. Relative mRNA levels were calculated with normalization to the housekeeping gene *GAPDH*. (NB. GAPDH did not change after chemotherapy treatment as assessed in the RNAseq analysis).

**Table 7:**
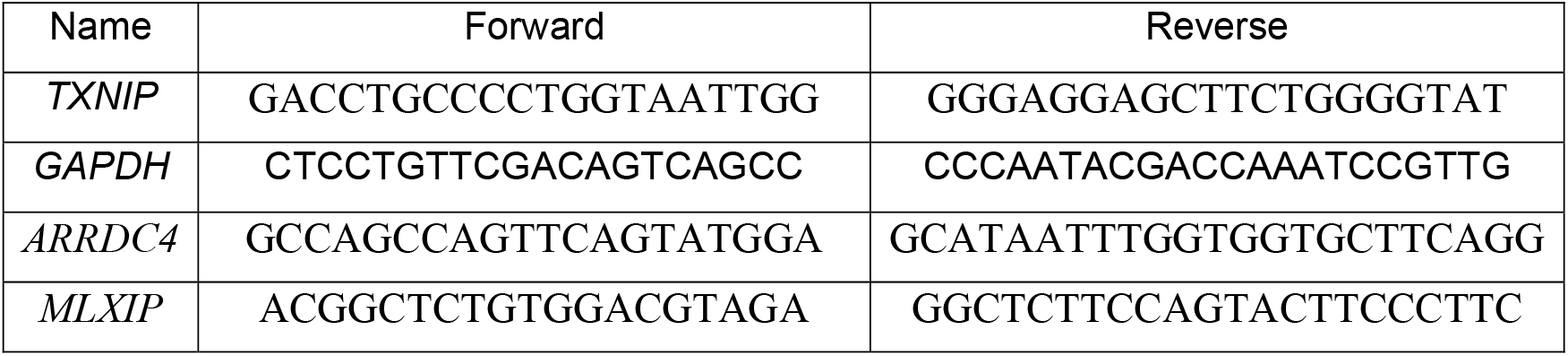
Primers for qRT-PCR.

### Chromatin Immunoprecipitation-Quantitative Polymerase Chain Reaction (ChIP-PCR)

DLD1 cells were seeded in 75cm^2^ flasks (∼40% confluency). Overnight, cells were replaced with fresh media for 1 h and then either treated or not treated with oxaliplatin/ NAC as indicated in the Figures. After 48 h, cells were cross-linked with 1% formaldehyde and quenched by glycine. Chromatin extraction was performed using the Chromatin Extraction Kit (ab117152) followed by sonication. Equal amount of chromatin was incubated overnight at 4°C with 2 µg of anti-MondoA (Proteintech, 13614-1-AP) or IgG (Cell Signaling Technology, 2729). ChIP pull-down assays were performed using the ChIP Kit Magnetic One-Step (ab 156907) according to the manufacturers’ instructions. Recovered DNA was quantified by qRT-PCR using primers specific for TXNIP promoter region (forward-CACAGCGATCTCACTGATTG; reverse-GTTAGTTTCAAGCAGGAGGC) under the following conditions: 40 cycles of denaturation at 95 °C for 15 s and annealing at 56 °C for 20 s, followed by extension at 72 °C for 40 s. Specificity of the PCR product was assessed by Sanger sequencing.

### Spheroid formation Assay

Spheroid culture was performed using suspensions of cells with at least 90% viability. The spheroid formation was performed with 1,000 vital cells in 100 µl per well in a low-attached 96 well plate (Corning, 3474) under standard culture conditions. DLD1 spheroids were formed after 24 h of seeding. HCT15 spheroids were formed after 72 h of seeding. CellTiter-Glo® 3D cell viability reagents (Promega) were used to analyse spheroid viability as per manufacturer’s instrcutions. Three-dimensional cultures were treated with oxaliplatin and incubated for 48 hours.

### Patient-derived tumor organoids (PDTOs)

University College Hospital London (UCLH) provided us with colonic tissues from colorectal cancer patients in accordance with the guidelines of the European Network of Research Ethics Committee (EUREC) following European, national, and local law. HTA licence: 12055, REC reference: 15/YH/0311 as overarching biobank ethical approval. Informed consent forms were signed by all the participants in the study. Patient consent can be withdrawn at any time, resulting in the prompt disposal of the tissue and any derived material.

CRC cells were isolated as described by Sato et al^72^. Briefly, specimens were washed with 10ml PBS and then cut into small pieces (1-2 mm) with 10ml of digestion buffer (Suppl Table 5). Tissue and digestion buffer were transferred to a gentleMACS C Tube (run protocol 37C_h_TDK_1) (Miltenyi Biotec, 130-096-334) and incubated at 37 °C for 1 hour. Supernatant was aspirated after samples were filtered through 100 µm strainers (732-2759) into 50 ml tube, and centrifuged at 800xg for 2 mins. After incubating with ACK lysis buffer (A1049201) at room temperature for 5 mins, samples were washed with PBS twice. Cell pellet was resuspended in appropriate volume of Matrigel and 40 µL organoid: Matrigel droplets were plated into a 6-well plate.

After incubation at 37°C for 10-20 min, 2 ml of complete medium (Suppl Table 5) supplemented with the ROCK Inhibitor Y-27632 (10 µM, 72302) were added in each well. Medium was changed twice a week until ready for passage. For qPCR and western blot analyses, organoids were seeded in 6 well plate and collected after drug treatments indicated.

### ROS measurement

ROS level in cells was detected using DHE (Dihydroethidium) Assay Kit—Reactive Oxygen Species (Abcam, ab236206). Around 1 × 10^5^ cells were added to V-bottom plate. 130 µL ROS staining buffer and then 100 µL Cell-Based Assay Buffer were used according to manufactures’ guides. The fluorescence was measured using an excitation wavelength between 480-520 nm and an emission wavelength between 570-600 nm.

### CRISPR-CAS9 genome engineering

MondoA, TXNIP and GDF15 knockouts in cells and organoids were carried using the CRISPR/Cas9 system and the Edit-R CRISPR/Cas9 gene engineering protocol (Horizon). Guide RNAs for TXNIP (Edit-R CRISPR (knockout) Human TXNIP crRNA, Catalog ID:CM-010814-01-0002), GDF15 (Edit-R CRISPR (knockout) Human GDF15 crRNA, Catalog ID:CM-019875-01-0002), and MondoA (Edit-R CRISPR (knockout) Human MLXIP crRNA, Catalog ID:CM-008976-01-0002) were purchased from Horizon.

Cells were transfected in a 6-well plate with crRNA: tracrRNA transfection complex and Cas9 mRNA, using DharmaFECT Duo Transfection Reagent (Horizon, T-2010-02) (Suppl Table 6). After 48 h, a BD Aria Fusion cell sorter was used to sort GFP-positive single cells into 96-well plates. To measure TXNIP and GDF15 levels, each clone was expanded for 3–6 weeks. The following knockout clones were chosen: Three TXNIP knockout clones, three MondoA knockout clones, and four GDF15 knockout clones. A heterogenous knockout cell line was generated by combining knockout clones of each gene and their functional evaluation was performed. Stabilities of the knockouts were checked every five passages using PCR and western blot analysis.

The neon® Transfection System (Thermo Fisher Scientific, MPK5000) was used for CRISPR Editing of organoids. 1 × 10^5^ organoids were trypsinized and single cells were resuspended in 7.5 µL of Resuspension Buffer R per electroporation condition, then 7.5 µL of RNP Complex Mix was added (Suppl Table 6). The mixture was electroplated as shown in Suppl Table 6. Immediately after electroporation, organoids were seeded onto a 24-well prewarmed plate. Complete medium was changed every 2 days and genome editing efficiency was assessed using PCR and western blot analysis.

### Mass Spectrometry

DLD-1 cells were seeded with a density around 70-80% in 6-well plates. On second day, cells were washed with PBS and replaced with 2 ml of FBS-free media (RPMI+1% penicillin/streptomycin +1% Glutamin). After 48 hrs (day 4), supernatants from cell culture were collected, centrifuged (300 g/ 5 min) to get remove debris, followed by adding cold acetone at a ratio of 1:3. The mix was shaken thoroughly and stored at −20°C overnight. Protein pellets were collected after a centrifugation at 10000 g for 15 min). Keep the pellets in −80°C freezer for storage till mass spectrometry analysis. Each protein pellet was resuspended in 20 µl of 8M urea, followed by adding NuPAGE™ LDS Sample Buffer (4X) (ThermoFisher). The mixture was kept at 90°C for 5min and loaded into a 10% Bis-Tris gel, resolved for about 1cm (80 volts; 63 mA; 8 watts) before being stained with Imperial protein stain (ThermoFisher). After de-staining to remove the background, the whole section was excised and followed by an in-gel trypsin digestion overnight at 37°C. 500 µg of TMTpro reagents (ThermoFisher) were added to the peptides (50 µg) along with acetonitrile and then incubated at room temperature for 1 h. After the labelling efficiency was checked out, the reaction was quenched with hydroxylamine to a final concentration of 0.3% (v/v) for 15min and all individual tags were combined as one. The sample was vacuum centrifuged to near dryness and subjected to C18 solid-phase extraction (SPE, Sep-Pak) for a clean-up.

MS data were collected using Orbitrap Fusion Lumos mass spectrometers. Orbitrap Fusion Lumos mass spectrometer was equipped with an Ultimate 3000 RSLC nano pump. Raw mass spectrometry data were processed into peak list files within Proteome Discoverer (ThermoScientific v2.5). Processed data were then searched using Sequest search engine embedded in Proteome Discoverer v2.5 against the reviewed Swissprot Homo Sapiens database downloaded from Uniprot (http://www.uniprot.org/uniprot/).

### Proteome profiler antibody array

Human (R&D Systems, ARY005B) cytokine arrays were used. Cells were seeded at 4 × 10^5^/well in 6-well plate. Next day, the cells were replaced with flesh media with or without indicated drug. Tumor-conditioned medium (TCM) was collected after 48 h of treatment. 0.5ml of TCM was added to membrane and soluble Proteome was analysed following manufacturer’s instructions.

### Enzyme-Linked Immunosorbent Assay (ELISA)

ELISAs for GDF15 (DGD150), IL-1β (DY201-05), and IFNγ(DY285B-05) were purchased from Biotechne and carried out as per the manufacturer’s instructions. Plates were read on a CLARIOstar instrument at 450 nm, being corrected against 570 nm, and analysed using MARS software and excel. The concentration of each sample was calculated using a standard curve.

### Immunofluorescence staining

5 × 10^3^ DLD1 cells were plated into 35 mm glass bottom dishes. After 24 h, cells were treated with 10 µM oxaliplatin. 48 hours post treatment, cells were rinsed with PBS, fixed for 20 min with 4% PFA, rinsed with PBS, permeabilized 10 min with 0.1% Triton-X100, rinsed with TBS-T. Subsequent labelling, imaging, and image analysis steps were as previously described^73^.

### Generation of CRISPRa Constructs

#### dCas9-VPR

The 10XUAS-dCas9-VPR constructs have been previously described^74^. Instructions are available at Addgene (https://www.addgene.org/78897/).

#### Transfection of stable dCas9-VPR expressing cell lines with synthetic guide RNAs

Cells were seeded in 6-well plates and cultured >50% confluency. Culture media was replaced with 1.6ml of fresh media before transfection. Transfection reagents were prepared in two separated tubes (A and B): Tube A (195 µl Serum/antibiotic-free media and 5 µl 10 µM guide RNA mix) (Table 8) and Tube B (195 µl Serum/antibiotic-free media and 5 µl DharmaFECT reagent 1). Tubes A and B were mixed thoroughly and incubated at room temperature for 20 min before being added to the cells.

**Table 8:** Sequence of crRNA oligonucleotides.

| Reagent or Resource | Source | Identifier |
| --- | --- | --- |
| CRISPRmod CRISPRa (activation)<br>Human MLXIP Synthetic crRNA<br>(SMARTpool) | Horizon | P-008976-01-0005 |
| CRISPRmod CRISPRa (activation)<br>Human TXNIP Synthetic crRNA<br>(SMARTpool) ) | Horizon | P-010814-01-0005 |
| CRISPRmod CRISPRa (activation)<br>Human GDF15 Synthetic crRNA<br>(SMARTpool) | Horizon | P-019875-01-0005 |
| CRISPRmod CRISPRa (activation)<br>Human MYC Synthetic crRNA<br>(SMARTpool) | Horizon | P-003282-01-0005 |
| CRISPRmod CRISPRa synthetic<br>crRNA non-targeting controls | Horizon | U-009500-10-05 |

### Immune cell isolation

Leucocyte cones were ordered from the National Health Service Blood and Transplant Service (NHSBTS) (The NHSBTS obtains informed consent from the donors and has internal ethical approval under the terms of their own HTA licence). Cells were mixed 1:1 with phosphate-buffered saline (PBS) and layered on Ficoll–Paque (GE Healthcare; 1714402). Cells were spun at 800 g for 30 min, with the brake off, and the PBMCs were taken from the buffy layer above the Ficoll–Paque. Naïve CD4 T cells were isolated from PBMCs using the MACS system as per manufacturer’s instructions (Miltenyi Biotech; 130-094-131. LS Columns; 130-042-401). Purity was checked using anti-CD4 and anti-CD45RA antibodies and seen to be > 95%. If purity was below 95%, the cells were disposed of.

### Flow cytometry

1-2 × 10^5^ cells were stained with a live/dead dye (ThermoFisher; L23102) in PBS for 10 min on ice in the dark, before being washed twice in FACS buffer (0.5% bovine serum albumin [Sigma; 05482] in PBS + 2 mM EDTA). Cells were then Fc blocked with Trustain (Biolegend; 422302) in FACS buffer for 10 min on ice in the dark. Cells were washed and then stained using a variety of antibodies ± secondary reagents described in Table 9, using concentrations recommended by the manufacturer, on ice for 30 min in the dark. Cells were washed and either read immediately or fixed using 1% PFA in FACS buffer and read within 3 days. Cells were read using a BD Accuri C6 Plus flow cytometer, with analysis carried out using BD Accuri C6 Plus software. All cells were gated as follows: (a) Forward scatter and side scatter (SSC) to exclude cellular debris (whilst also adjusting threshold), (b) live/dead (only live cells carried forward) and (c) SSC-A vs. SSC-H—only singlets carried forward. All MFIs were corrected against an appropriate isotype control. Intracellular flow cytometry was carried out using the intracellular fixation and permeabilization kit (ebioscience; 88-8824-00) according to manufacturer’s instructions.

**Table 9:** antibodies and reagents.

| Antibodies | Source | Identifier |
| --- | --- | --- |
| LIVE/DEAD™ Fixable Red Dead Cell Stain Kit | ThermoFisher | Cat# L23102 |
| Human TruStain FcX™ (Fc Receptor Blocking Solution) | Biolegend | Cat# 422302 |
| PE Mouse IgG1, κ Isotype Ctrl Antibody | Biolegend | Cat# 400112 |
| FITC Mouse IgG1, κ Isotype Ctrl (FC) Antibody | Biolegend | Cat# 400110 |
| PerCP Mouse IgG1, κ Isotype Ctrl Antibody | Biolegend | Cat# 400148 |
| FITC anti-human CD48 Antibody | Biolegend | Cat# 336706 |
| PerCP anti-human CD4 Antibody | Biolegend | Cat# 317432 |
| FITC anti-human CD3 Antibody | Biolegend | Cat# 317306 |
| PE anti-human CD8 Antibody | Biolegend | Cat# 344706 |
| PE anti-human FOXP3 Antibody | Biolegend | Cat# 320108 |
| PE anti-human CD45RA Antibody | Biolegend | Cat# 304108 |
| FOXP3 Fix/Perm Buffer Set | Biolegend | Cat# 421403 |

### Proliferation assays

96 well tissue culture stimulation plates were prepared the night before by adding 100 µl/well 1 µg/ml anti-CD3 (OKT3) in PBS. PBMCs were stained using an eFluor^TM^ 670 dye (65-0840-85; ebioscience) according to manufacturer’s instructions, and plated at 2×10^5^ cells in 100 µl. 100 µl of supernatant or other factors were added and cells were cultured for 4 days.

### Functional Treg assay

Anti-CD3 (OKT3) was plated at 1 µg/ml in PBS and incubated overnight at 4 °C. Supernatant was removed and 2×10^5^ / cell isolated naïve CD4 cells were added in the presence of 1 µg/ml anti-CD28 in the presence of NTC or GDF15 (a) supernatant +/-isotype control (10 µg/ml) or anti-CD48 (10 µg/ml). Cells were cultured at 37 °C for 4 days. On day 3, anti-CD3 was plated at 1 µg/ml in PBS and incubated overnight at 4 °C. Allogeneic PBMCs were isolated, stained with eFluor^TM^ 670 proliferation dye and plated at 1×10^5^ cells/ well. 1×10^5^ Tregs were added at a 1:1 ratio and the co-culture was run for 4 days. Cells were then harvested and stained with anti-CD3, anti-CD8 and anti-CD4 antibodies. The proliferation dye MFI in the responder population was normalized against matched cells stimulated in media alone.

### Establishment of oxaliplatin-resistant (OXAR) cell lines

Oxaliplatin-resistant cells (OXAR) cells were established by treatment with constant oxaliplatin concentration *in vitro*. Different oxaliplatin concentrations (50 µM for DLD1 and 25 µM for HCT15) were added to RPMI complete media. DLD1 and HCT15 cells were sub-cultured every 2 weeks. Finally, cell lines that capable of growing exponentially in RPMI with high concentrations of oxaliplatin were identified as drug resistant cell lines. The final tolerated drug concentrations are shown in Table 10. Experiments on resistant cell lines were performed after culturing in the medium without oxaliplatin for at least 2-3 weeks.

**Table 10.** The tolerated concentration of each resistant subline from oxaliplatin.

| Drug resistant cell lines | Drug concentration (IC50, ratio) |
| --- | --- |
| DLD1-OXAR | 109.20 µM (=6.2×IC50) |
| HCT15-OXAR | 36.45 µM (=5.4×IC50) |

### Cell viability Assay

The Deep Blue Cell Viability^TM^ Kit (BioLegend, 424701) was used to analyse cell chemotherapy-induced cytotoxicity. After cells were seeded into 96-well plates (5000 cells/well), oxaliplatin (Ebewe Pharma, Austria) was added to the wells in several doses for 48-72 hours. The plate was incubated at 37 °C for 3 hours following the addition of 1:10 volume ratio of Deep Blue Cell Viability™ reagent to each well. A CLARIOstar Plate Reader (Excitation: 530-570 nm, Emission = 590-620 nm) was used to detect the reduction of resazurin into resorufin and the OD value was used to calculate cell viability.

### Statistical analysis

All *in vitro* experiments were performed in three independent replicates for three times. All quantitative data are presented as mean ± standard error of the mean (SEM) and were analysed using GraphPad Prism 9.0. The means of the two datasets were compared using paired t-tests. One-way ANOVA was used to evaluate multiple independent groups. The chi-squared test was applied to compare categorical variables. Kaplan–Meier analyses were performed via the survival package. P-value<0.05 was considered as statistically significant.

## Acknowledgements and Funding

This work was supported by China Scholarship Council Awards (No.201806010012 to JD, No.202006940028 to TP); CRUK Early Detection and Diagnosis Committee (Project grant, C1519/A27375). RB is supported by MR/R000026/1 and UCLH/UCL BRC who received a proportion of funding from the Department of Health’s NIHR Biomedical Research Centres funding scheme. LD is supported by EU IMI2 IMMUCAN (Grant agreement number 821558). GA and JV are supported by CRUK Early Detection and Diagnosis Committee Project grant (C7675/A29313). ZA was supported by the KCL Breast Cancer Now Research Unit (grant KCL-Q2-Y5). KN was supported by Cancer Research UK Clinical Training Fellowship (Award number 176885). CG is supported by CRUK City of London Centre (CTRQQR-2021\100004). RE is jointly supported by a Global Pharmaceutical Development Science Fellowship and Training grant to KCL and CRUK City of London Centre RadNET grant.

## Authors’ contributions

JD and TP have contributed equally to this work. TN and RB are the corresponding authors to this work. Conceptualization, resources, supervision and project administration were carried out by TN, JD and RB; Methodology, software, formal analysis, investigation, data curation and visualization were performed by JD, TP, RB. Writing – original draft by JD, RB. Writing – review & editing were done by TN, JD, RB and TP. Single cell RNA sequencing analyses - YH, ZH and ST. Patient derived organoids establishment: M R-J, PV, CJT, JD, KN, CADCG, CM. Mass spectrometry analyses: XY, JD, TP. Patient sample collection and IHC staining: XZ, GL. TCGA data analyses: ZL, LL, YC. ChIP-PCR analysis: GA, JD. Q-PCR analysis: JD, TP, LD. CRISPR-activation cell model establishment: JM, JD. MY, JCP, JMV and GW reviewed and edited the manuscript. All authors reviewed and approved the final manuscript.

## Ethics approval and consent to participate

The study was performed in accordance with the principles of the Declaration of Helsinki.

## Consent for publication

We have obtained consent to publish from the participant to report individual patient data.

## Availability of data and material

The RNAseq data will be available to the public through the GEO portal (currently in process). Other datasets generated during and/or analysed during the current study are available from the corresponding author on reasonable request.

---

**Figure S1.**
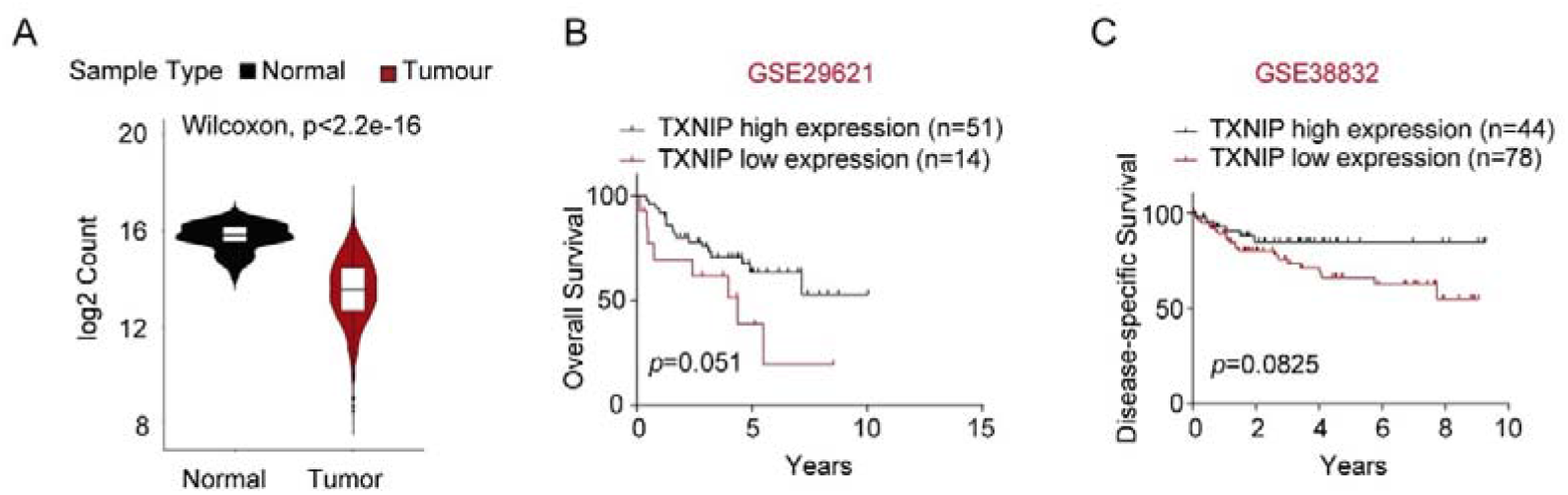
TXNIP expression is lower in colorectal cancer samples compared to normal tissues. (A) Analysis of The Cancer Genomic Atlas (TCGA) Colon Adenocarcinoma (COAD) database. Comparative analysis of TXNIP transcript expression between adjacent normal tissue and cancer tissues. (B-C) Kaplan-Meier analysis of overall survival (B) and distant metastasis-free survival (C) in CRC patients with different TXNIP mRNA expression levels. Wilcoxon rank-sum test p value indicated.

**Figure S2.**
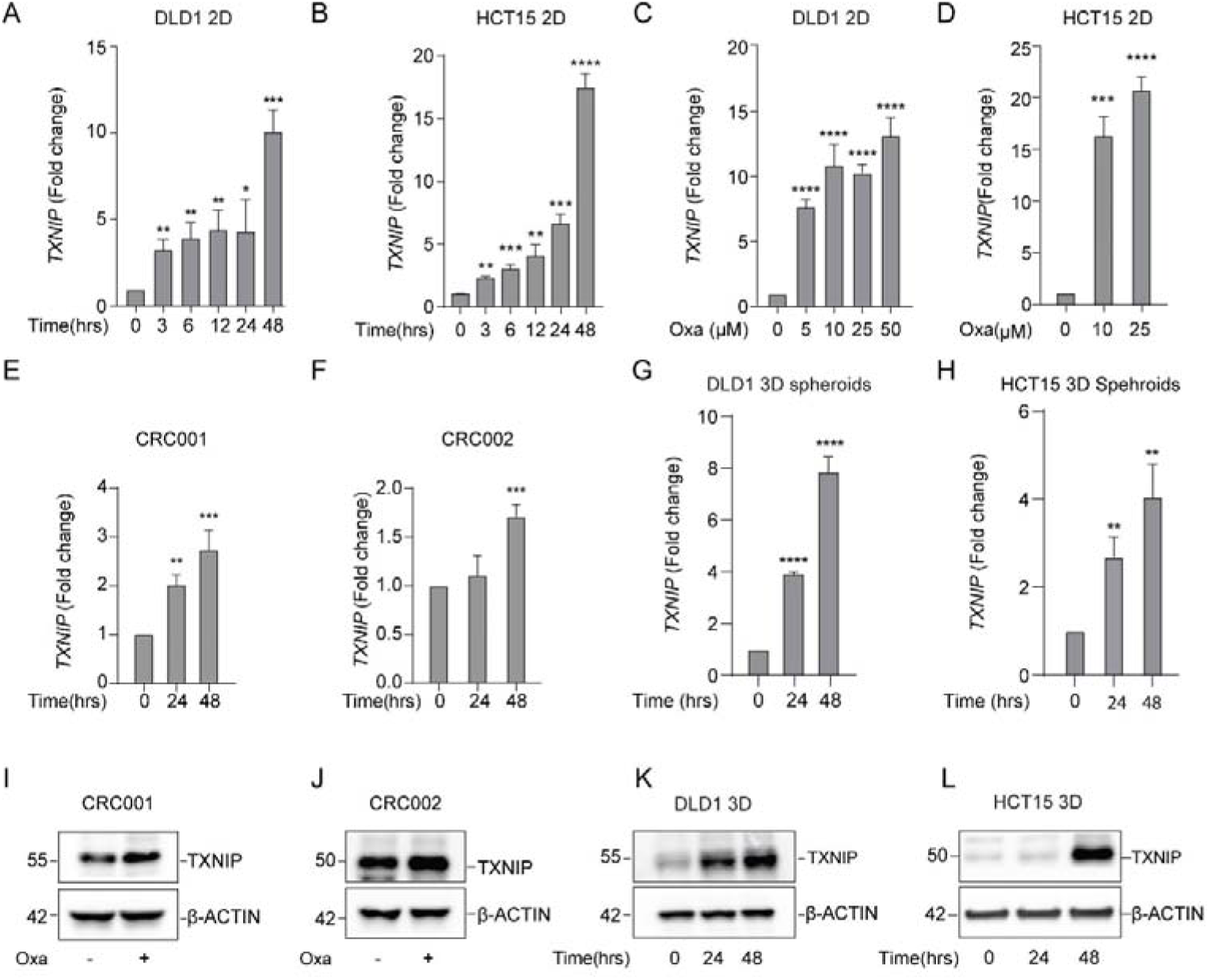
TXNIP expression is induced by oxaliplatin in different CRC models. (A-B) Assessment of *TXNIP* mRNA expression in DLD1 cells (A) or HCT15 cells (B) treated with oxaliplatin by q-RT-PCR analysis. Cells were treated with 10µM oxaliplatin and harvested at indicated time points. (C-D) q-RT-PCR analysis of *TXNIP* mRNA in DLD1 cells (C) or HCT15 cells (D) treated with oxaliplatin for 48h at indicated concentrations. (E-F) q-RT-PCR analysis of *TXNIP* mRNA in two different PDTOs treated with 10µm oxaliplatin for indicated time periods. (G-H) q-RT-PCR analysis of *TXNIP* mRNA in DLD1 (G) or HCT15 (H) spheroids treated with 10µm oxaliplatin for indicated time periods. (I-J) Western blotting analyses of TXNIP post oxaliplatin treatment (10µm) in two different PDTOs for 48h. (K-L) Western blotting of TXNIP in DLD1 (K) or HCT15 (L) spheroids treated with 10µm oxaliplatin for 48h. Results shown are representative of three independent experiments. All values were expressed as mean ± SEM. *p<0.1, **p<0.01, ***p < 0.001, ****p < 0.0001, vs. Control.

**Figure S3.**
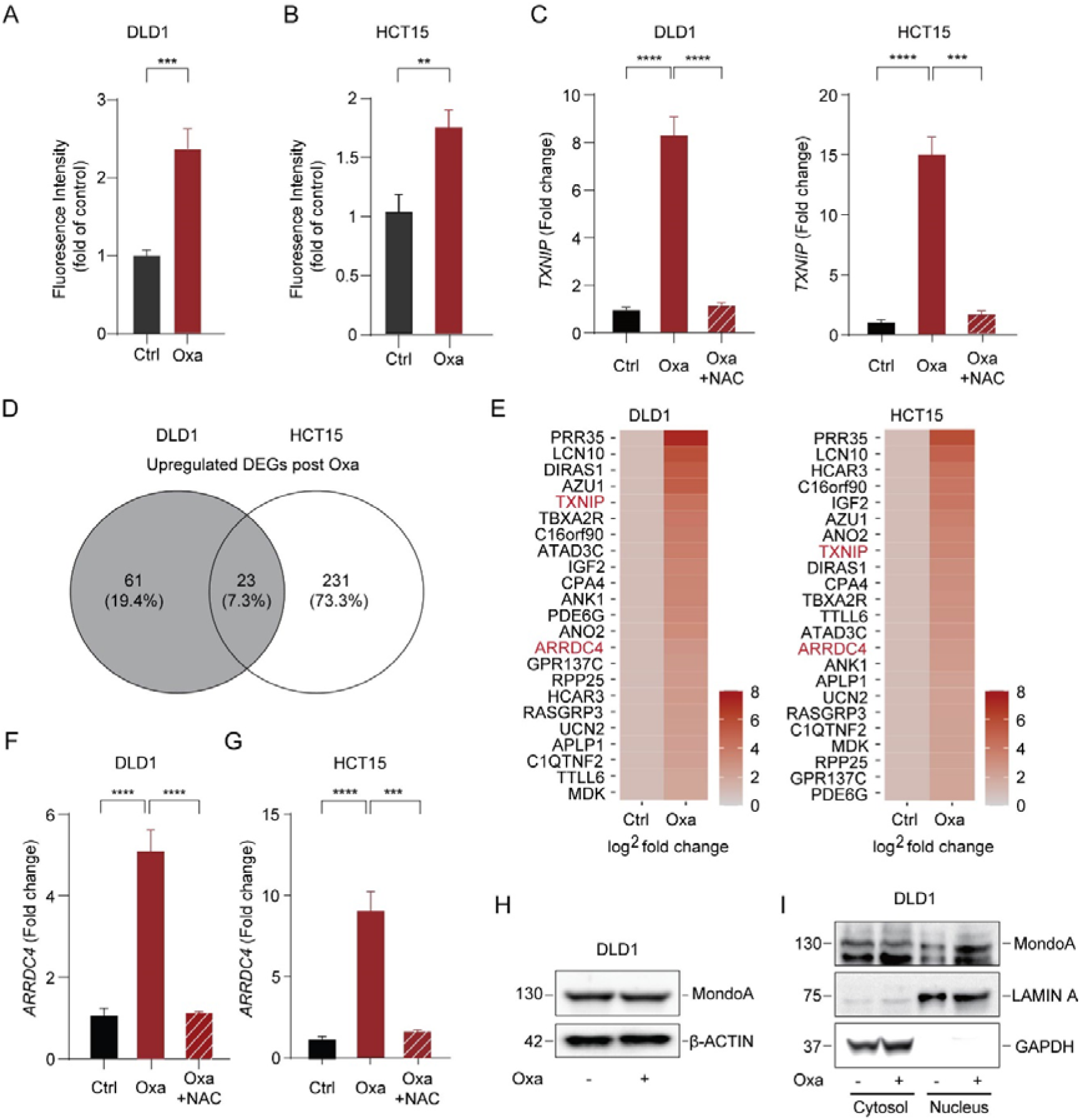
ROS drive the induction of TXNIP by inducing MondoA activity. (A-B) DLD1 cells (A) and HCT15 cells (B) were treated with 10µm oxaliplatin with ROS measured at 48h. (C) qRT-PCR analysis of *TXNIP* mRNA in DLD1 cells (left panel) or HCT15 cells (right panel) treated with N-acetyl-L-cysteine (NAC) (1.25mM) or oxaliplatin (10µm), or combinational treatment, for 48h. (D) Overlapping DEGs (>4-fold change; Padj<0.05) from live DLD1 and HCT15 cells, after 48h of 10µm oxaliplatin treatment, as determined by RNA sequencing. (E) Heatmap showing 23 overlapping transcripts from D, in DLD1 cells (left panel) and HCT15 cells (right panel). (F-G) qRT-PCR analysis of *ARRDC4* mRNA in DLD1 cells (F) and HCT15 cells (G) treated with with NAC (1.25mM) or oxaliplatin (10µm), or combinational treatment, for 48h. (H) Immunoblot analysis of MondoA expression in DLD1 cells after 10µm oxaliplatin treatment for 48h. (I) Effects of oxaliplatin treatment (10µm for 48h) on subcellular localization of MondoA assessed by cell fractionation and immunoblotting, in DLD1 cells. LAMIN A - a nuclear marker, GAPDH - a cytoplasmic marker. Results shown are representative of three independent experiments. All values were expressed as mean ± SEM. **p<0.01, ***p < 0.001, ****p < 0.0001, vs. Control

**Figure S4.**
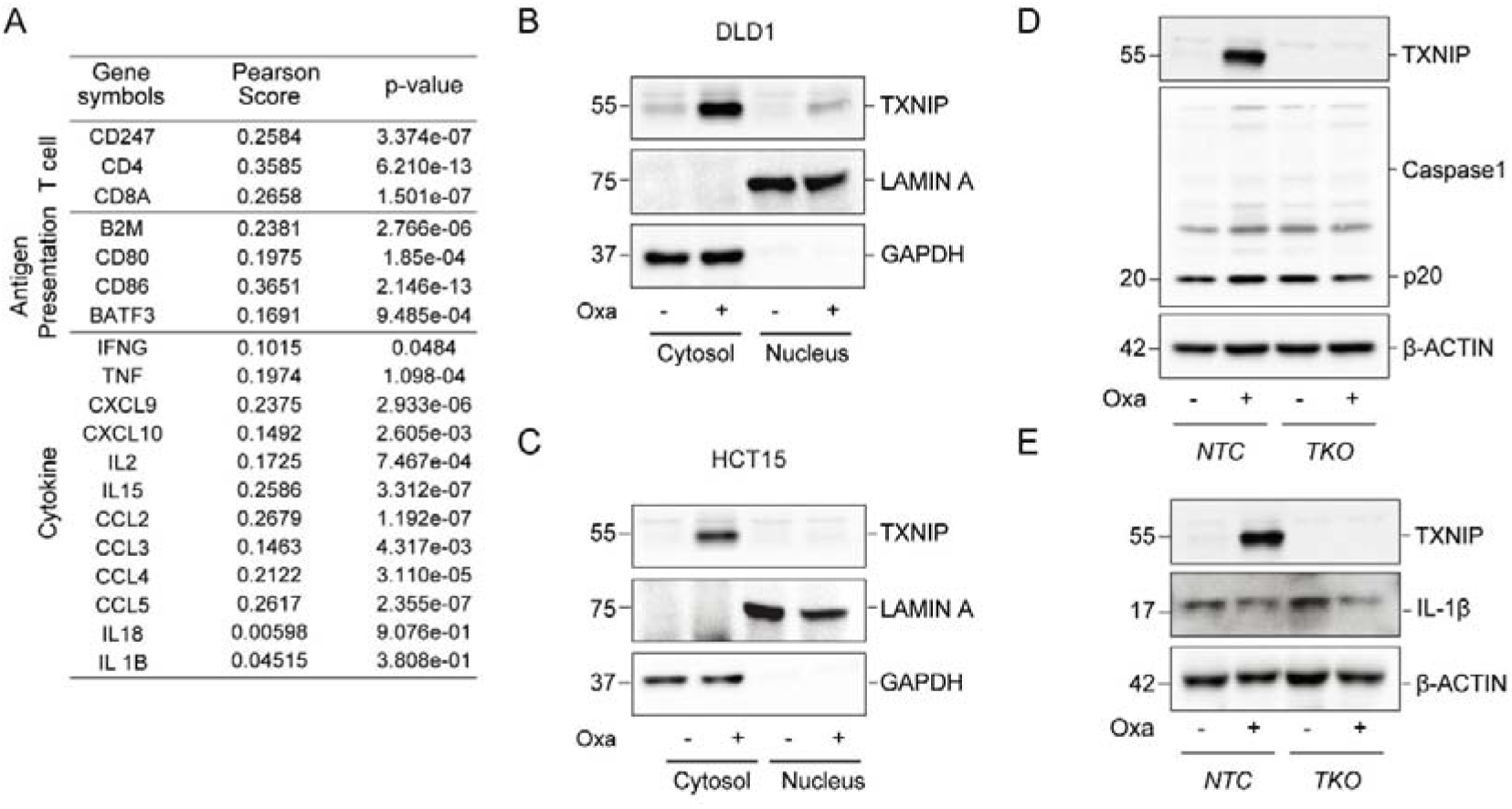
TXNIP is associated with immune activation, which is independent of inflammasome activity. (A) Pearson correlation coefficient scores and p values showing the relationship between *TXNIP* transcript expression and different immune marker transcript expression; including T cell markers (*CD247, CD4, CD8A*), antigen presentation markers (*B2M, CD80, CD86, BATF3*) and cytokines (*IFNG, TNF, CXCL9, CXCL10, IL2, IL15, CCL2, CCL3, CCL4, CCL5, IL18, IL1B*) from the TCGA COAD dataset. (B-C) Effects of oxaliplatin (10µm for 48h) on subcellular localization of TXNIP assessed by cell fractionation and immunoblotting in DLD1 cells (B) and HCT15 cells (C). (D-E) Immunoblot analysis of cleaved caspase 1(p20) (D) and IL-1β (E) in control (NTC) and TXNIP-KO (TKO) DLD1 cells with/ without 10µm oxaliplatin treatment for 48h. Results shown are representative of three independent experiments.

**Figure S5.**
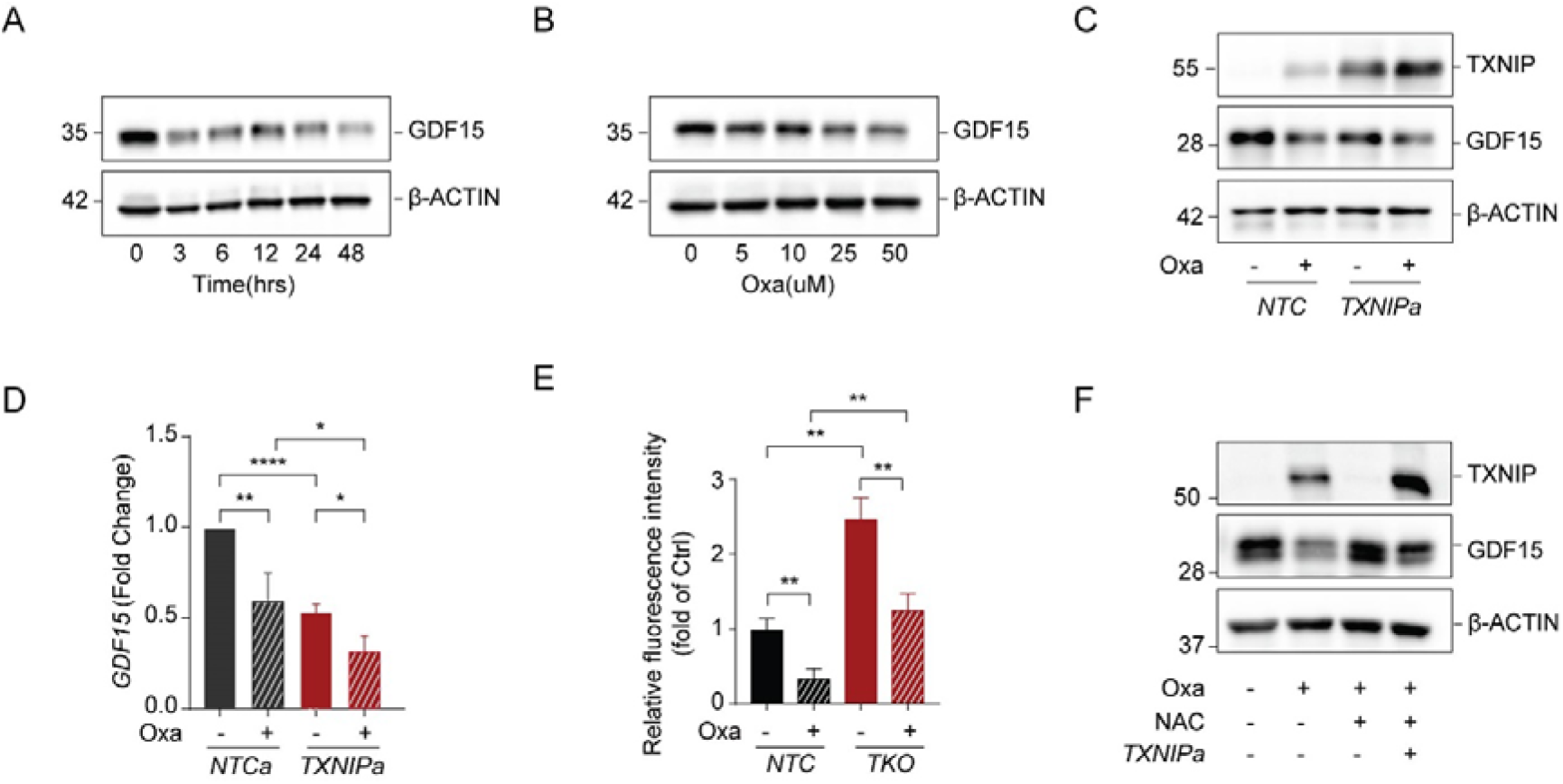
Oxaliplatin treatment and TXNIP suppress GDF15 expression. (A-B) Immunoblotting of GDF15 in DLD1 cells after treatment with 10µm oxaliplatin at indicated time points. (A); after treatment of different dosages of oxaliplatin for 48 hours (B). (C-D) Immunoblotting of TXNIP and GDF15 in control (NTC) and TXNIP-overexpressing (TXNIPa) DLD1 cells with or without 10µm oxaliplatin treatment for 48h (C); pooled densiometric data from C (D). Standard error bars are shown n=3. (E) Quantitation of immunofluorescence from Figure 3I (GDF15 levels relative to cell area) from 3 independent experiments. (F) Immunoblotting of TXNIP and GDF15 in TXNIPa or NTC cells treated with oxaliplatin (10µm) or combined treatment with oxaliplatin and NAC (1.25mM) for 48h. Results shown are representative of three independent experiments. All values were expressed as mean ± SEM. *p<0.1, **p<0.01, ****p < 0.0001, vs. Control.

**Figure S6.**
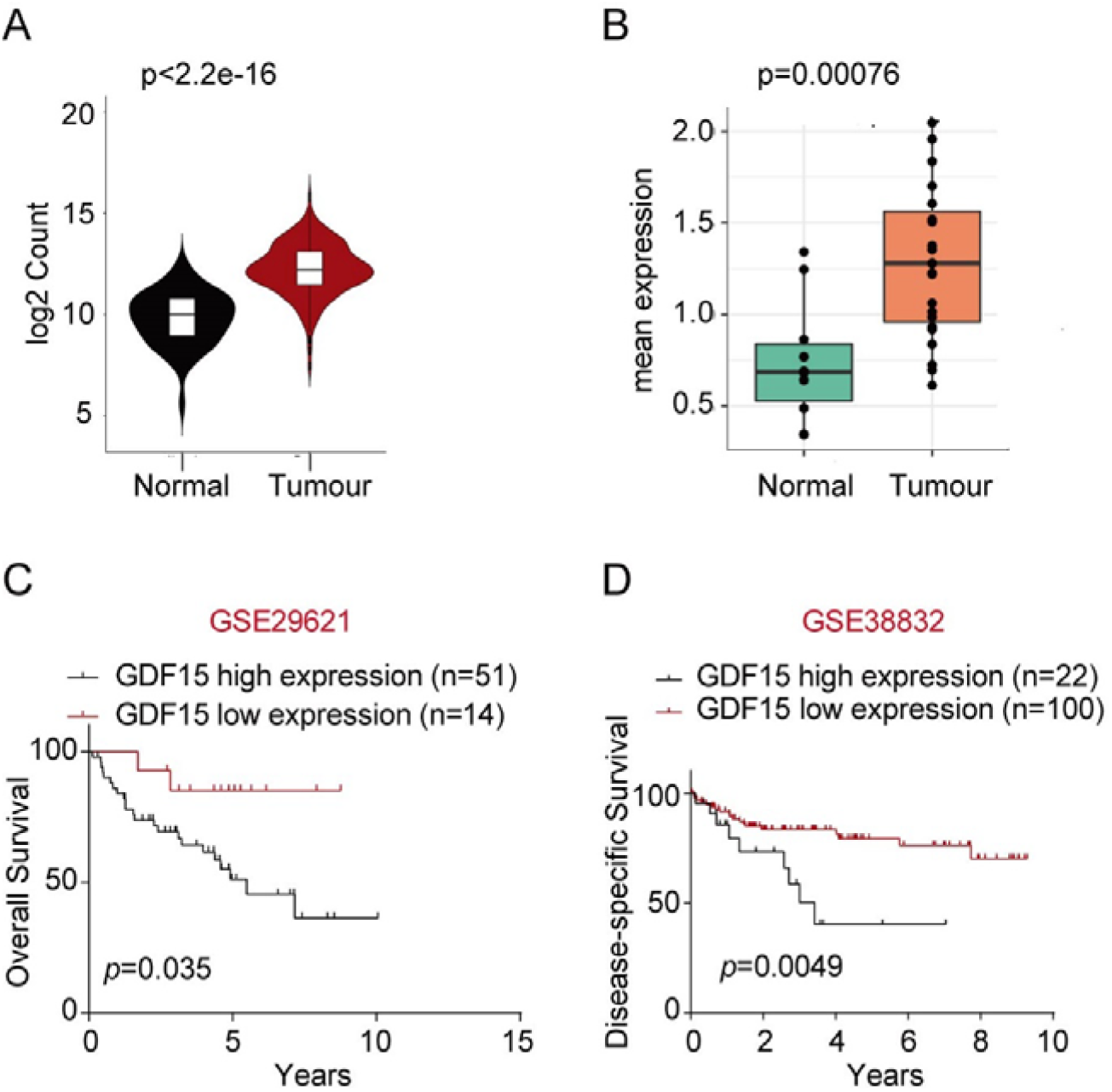
GDF15 expression is higher in colorectal cancer samples compared to normal tissues. (A) Analysis of The Cancer Genomic Atlas (TCGA) Colon Adenocarcinoma (COAD) database. Comparative analysis of expression of GDF15 between adjacent normal tissue and cancer tissues. Wilcoxon rank-sum test p value indicated. (B) GDF15 transcript expression in single epithelial cells derived from matched primary CRC tumors and adjacent normal colon (n=10 pairs). (C-D) Kaplan-Meier analysis of overall survival (C) and distant metastasis-free survival (D) in CRC patients with different GDF15 mRNA expression levels.

**Figure S7.**
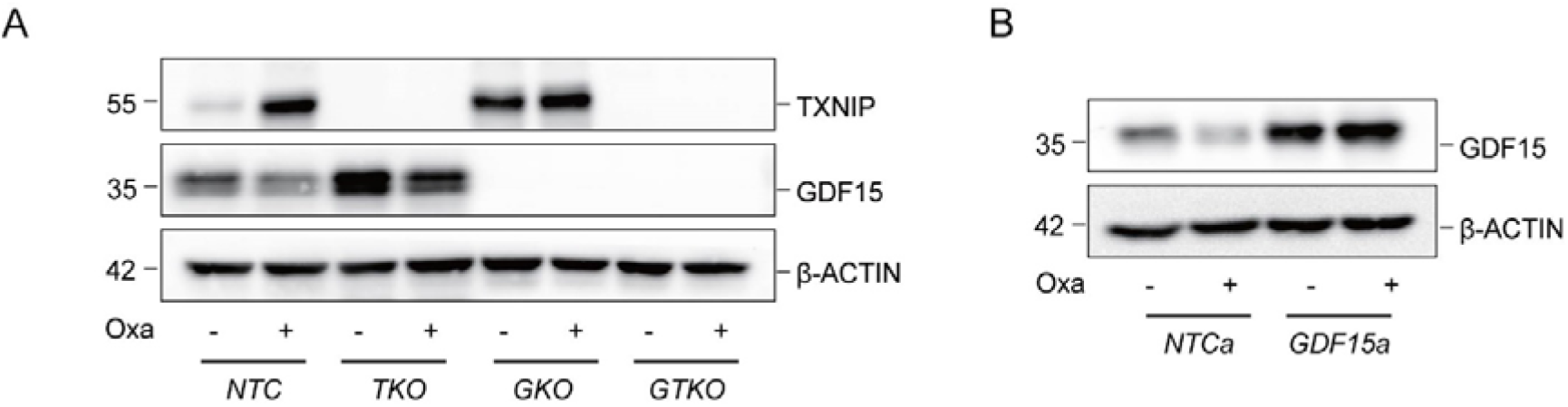
Establishment of knock-out and over-expressing DLD1 cell models. (A) Immunoblot of TXNIP and GDF15 expression in NTC, GDF15 knockout (GKO), TKO, GDF15 and TXNIP knockout (GTKO) DLD1 cell lines after 48h of oxaliplatin treatment (10µm). (B) Immunoblot of GDF15 expression in GDF15-CRISPRa (GDF15a) DLD1 cell line in the presence of 10µm oxaliplatin for 48h. Results shown are representative of three independent experiments.

**Figure S8.**
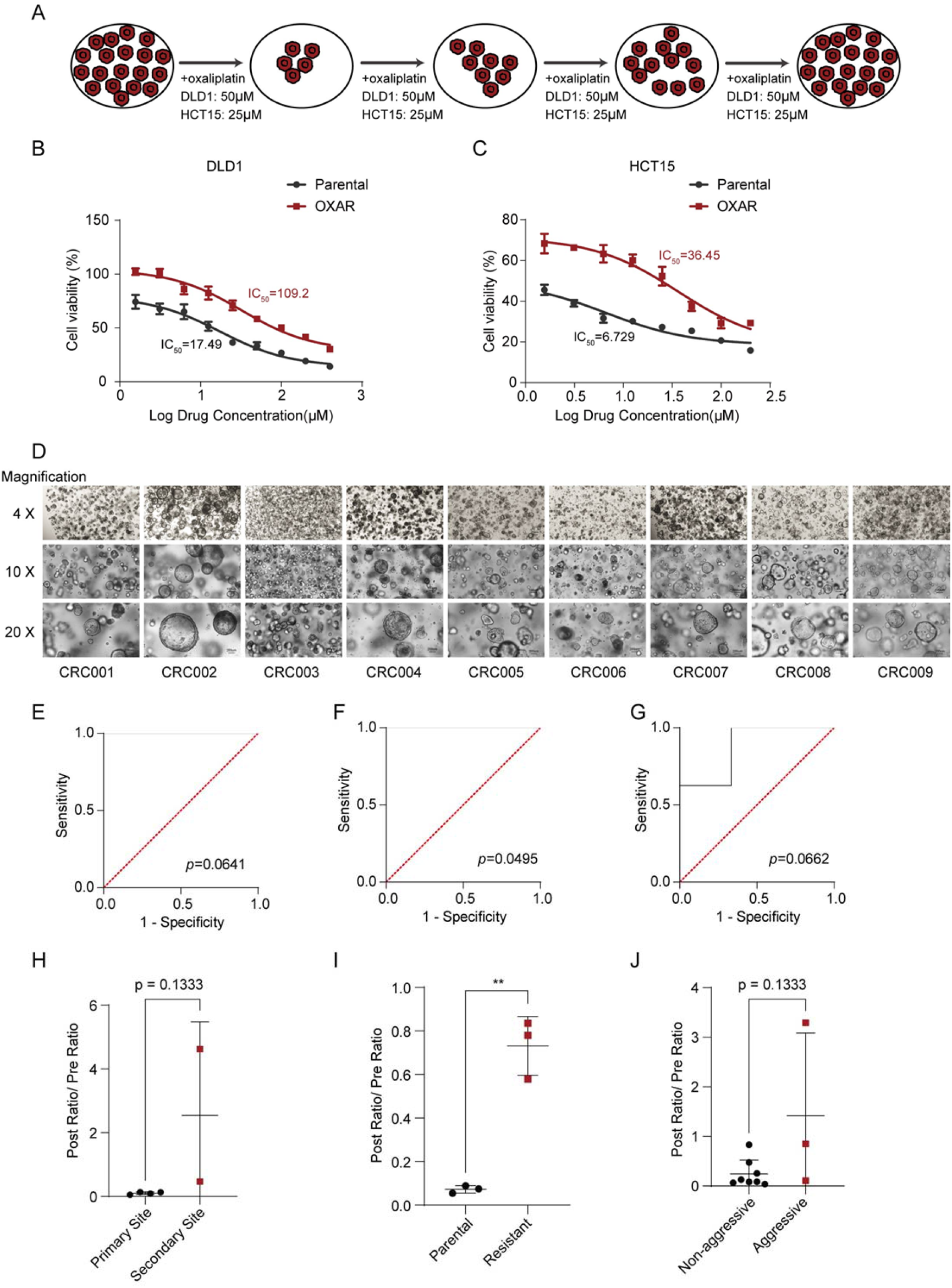
Establishment of oxaliplatin-resistant cell lines and patient-derived tumor organoids. (A) A schematic model showing the process by which oxaliplatin-resistant CRC cells were generated. (B-C) IC50 values of oxaliplatin in oxaliplatin-resistant cells (OXAR) and their parental cells. DLD1 and DLD1-OXAR (B); HCT15 and HCT15-OXAR (C). (D) Bright field images of different organoids at different magnifications. (E-G) Receiver operating characteristic (ROC) curves showing area under the curve and p values for the use of GDF15/TXNIP ratio in predicting origin of cell line (E; primary [n=4] or secondary [n=2]), sensitivity to oxaliplatin (F; parental [n=3] or resistant [n=3]), aggression of tumour (G; non-aggressive [n=8] or non-aggressive [n=3]). (H-J) Post-treatment GDF15/TXNIP ratio divided by pre-treatment GDF15/TXNIP ratio for primary or secondary cell line source (H), parental or resistant cell line (I), or aggression of fresh primary tumour (J). ** p<0.01 using unpaired t test. H and J tested using Mann-Whitney.

